# Knock-down of a regulatory barcode shifts macrophage polarization destination from M1 to M2 and increases pathogen burden upon *S. aureus* infection

**DOI:** 10.1101/2021.10.19.464946

**Authors:** Sathyabaarathi Ravichandran, Bharat Bhatt, Awantika Shah, Debajyoti Das, Kithiganahalli Narayanaswamy Balaji, Nagasuma Chandra

## Abstract

Macrophages are driven to form distinct functional phenotypes in response to different immunological stimuli, in a process widely referred to as macrophage polarization. Transcriptional regulators that guide macrophage polarization in response to a given trigger remain largely unknown. In this study, we interrogate the programmable landscape in macrophages to find regulatory panels that determine the precise polarization state that a macrophage is driven to. Towards this, we configure an integrative network analysis pipeline that utilizes macrophage transcriptomes in response to 28 distinct stimuli and reconstructs contextualized human gene regulatory networks, and identifies epicentres of perturbations in each case. We find that these contextualized regulatory networks form a spectrum of thirteen distinct clusters with M1 and M2 at the two ends. Using our computational pipeline, we identify combinatorial panels of epicentric regulatory factors (RFs) for each polarization state. We demonstrate that a set of three RFs i.e., *CEBPB*, *NFE2L2* and *BCL3*, is sufficient to change the polarization destination from M1 to M2. siRNA knockdown of the 3-RF set in THP1 derived M0 cells, despite exposure to an M1 stimulant, significantly attenuated the shift to M1 phenotype, and instead increased the expression of M2 markers. Single knockdown of each RF also showed a similar trend. The siRNA-mediated knockdown of the 3-RF set rendered the macrophages hyper-susceptible to *Staphylococcus aureus* infection, demonstrating the importance of these factors in modulating immune responses. Overall, our results provide insights into the transcriptional mechanisms underlying macrophage polarization and identify key regulatory factors that may be targeted to modulate immune responses.

## Background

Macrophages, the key components of the innate immune system, are responsible for processes ranging from the maintenance of healthy tissues, phagocytic clearance of microbes, to tissue repair and remodelling (Aderem, 2003; Greenberg and Grinstein, 2002; Mosser et al., 2021; Murray and Wynn, 2011). They do most of these in a bid to restore normal physiology and act as a bridge between innate and adaptive immunity. In response to a range of stimuli, including pathogen insults, macrophages are directed to distinct functional phenotypes in a process widely referred to as macrophage polarization (Benoit et al., 2008; Murray and Wynn, 2011). The most well-described phenotypes are the M1 and M2 polarization states, where M1 macrophages are pro-inflammatory in nature with microbicidal activity, while M2 is associated with an anti-inflammatory response. Stimulation of macrophages with lipopolysaccharides (LPS) or interferon-gamma (IFNγ) typically leads to the M1 state, whereas treatment with interleukins (IL4, IL13) polarize macrophages into the M2 state (Mills et al., 2000; Murray, 2017). M2 macrophages are further sub-categorised into M2a, M2b, M2c and M2d depending on the activation triggers and the resulting transcriptional changes (Mantovani et al., 2004; Rőszer, 2015; Yao et al., 2019). However, this model is said to be oversimplified as it fails to capture the complexity exhibited by different macrophage states in an *in vivo* scenario where the interaction between different cytokines, interleukins, and other effector molecules define the final polarized state (Atri et al., 2018; Martinez and Gordon, 2014). A recent study on the transcriptomes of human monocyte-derived macrophages treated with a variety of physiological stimulants identified nine different polarization states, which better fit the description of a spectrum of states rather than a classical binary M1/M2 axis (Xue et al., 2014). The different states in the polarization spectrum present complex transcriptional profiles as seen from altered gene expression patterns in several hundred genes. The importance of transcriptional regulation in defining macrophage polarization has been brought out by several studies (Gerrick et al., 2018; Hörhold et al., 2020; Palma et al., 2020; Xue et al., 2014; Zhao and Popel, 2021). *In vitro* and *ex-vivo* studies have shown complete reversal of macrophage polarization states depending upon the molecules (pathogen-associated molecular patterns (PAMPs), host-derived damage-associated molecular patterns (DAMPs) or cytokines, etc) present in the microenvironment.

Several studies have shown that activation of transcription factors (TFs) such as signal transducers and activators of transcription (STAT1, STAT3, STAT4 and STAT6), nuclear transcription factor-κB (NF-kB), NF-E2-related factors (NRF1 and NRF2), kruppel-like factors (KLF4 and KLF6), CCAAT-enhancer-binding proteins (C/EBP-α and C/EBP-β) and peroxisome proliferator-activated receptors (PPAR-α, PPAR-δ and PPAR-γ) are associated with macrophage polarization (Lawrence and Natoli, 2011; Li et al., 2018; Tugal et al., 2013). But, the available information is insufficient to predict the polarization state that will result in a given condition. A possible reason is that a set of TFs together defines the state rather than individual TFs by themselves. While the triggers that lead to different polarization states are known, the specific set of TFs that nucleate the change or serve as control points to guide macrophage polarization is not known.

The precise polarization state attained by the macrophages in response to pathogenic triggers have a direct bearing on the outcome of the disease (Benoit et al., 2008; Koziel et al., 2009; Pidwill et al., 2020). In response to bacterial pathogens such as *Staphylococcus aureus*, macrophages induce the transcriptional activity of pro-inflammatory genes, activating the M1 program. The M1 phenotype is known to have an activated microbicidal machinery that is utilized in clearing such acute infections. However, prolonged or excessive inflammatory response by M1 macrophages can be pathological as it results in tissue damage and destruction. M2 macrophages produce anti-inflammatory mediators and resolve the inflammation (Koziel et al., 2009; Pidwill et al., 2020). Not surprisingly, the skewing of macrophage polarization is observed in several cases of severe bacterial infections, sepsis, tumour progression in cancers, and a range of other inflammatory diseases (Ardura et al., 2019; Hamilton and Tak, 2009; Sica and Mantovani, 2012; Zheng et al., 2018). Although our understanding of macrophage biology is rapidly increasing, the precise molecular controls and their associated processes involved in macrophage switching to different functional states are not well characterized. Identifying combinations of molecules that guide macrophage polarization will greatly facilitate a deeper understanding of how infections are contained, provide a basis to predict the precise phenotype that will be attained in different situations and serve as a foundation for precision diagnostics and treatment strategies focused on the macrophages.

In this work, we seek to interrogate the programmable landscape in macrophages to identify regulatory panels that govern the precise macrophage polarization state. Towards this, we configured a computational pipeline that uses an integrative graph-theoretical approach where differential transcriptomes of macrophages were mapped onto the knowledge-based genome-scale gene regulatory network (hGRN), computed connected sets of top-ranked perturbations, and identified regulatory barcodes defining different polarization states ultimately. We then experimentally evaluated specific RFs in the M1 regulatory barcode in a THP-1 derived macrophage cell line. We also probed the role of these specific RF sets in the *S.aureus* infected human monocyte-derived macrophages and cell-line infection model. Together, we validated genes in the M1 barcode and demonstrated that knocking down specific RF sets in the barcode attenuated the expression of M1 markers and increased the expression of M2 genes. We further showed that such a state switch leads to hyper-susceptibility to *S.aureus* infection.

## Materials and Methods

### Transcriptomes of macrophage polarization states

We retrieved transcriptomes of macrophages stimulated with a diverse set of stimulants (GSE46903) from NCBI-GEO (Barrett, 2020; Xue et al., 2014). Briefly, the authors of this study purified monocytes from peripheral blood mononuclear cells (PBMCs) of healthy controls and subjected them to granulocyte-macrophage colony-stimulating factor (GM-CSF) or macrophage-colony stimulating factor (M-CSF) for 72 hours (hrs) to generate baseline macrophages (Mo). Then, they treated the Mo with one of the twenty-eight different stimulants for different time points ranging from 30 mins to 72 hrs and generated transcriptomes from each stimulant and each time point. The stimulants spanned across the broad spectrum of pro and anti-inflammatory triggers to generate a wide range of polarization states. The triggers and their end time-points (n=145) considered for this analysis are shown in **Table S1**. We chose this experiment for analysis since it had the maximum number of stimulants (n=28) compared to other publicly available datasets. Such an experiment will allow us to capture stimulant centric regulator factors without the effect of technical and biological confounders. Further, to study the relevance of macrophage polarization in acute bacterial infections, transcriptomes of peripheral macrophages exposed to *S. aureus* were retrieved from NCBI-GEO (GSE13670) (Koziel et al., 2009).

Raw data from the respective transcriptome experiments were retrieved and pre-processed in a platform-specific manner using appropriate packages in R (Carvalho et al., 2019; Gautier et al., 2004). Probes below the detection limit in ≥ 80% of the samples in each dataset were filtered out, and multiple probes associated with the same gene were summarized by taking an average. We then performed differential analysis for the macrophages treated with stimulants compared to their untreated counterparts, using the limma package in R (Ritchie et al., 2015). Genes with fold change ≥ ±1.3 and a False Discovery Rate corrected q-value ≤0.05 computed using the Benjamini-Hochberg procedure (Benjamini and Hochberg, 1995) were considered to be statistically significant differentially expressed genes (DEGs).

### Reconstruction of human Gene Regulatory Network (hGRN)

We reconstructed a comprehensive knowledge-based human Gene Regulatory Network (hGRN), consisting of Regulatory Factors (RF) to Target Gene (TG) and RF to RF interactions. Here, RFs refer to transcription factors and cofactors. To achieve this, we curated the experimentally determined regulatory interactions (RF-TG, RF-RF) associated with human regulatory factors (Wingender et al., 2013) in this study. These interactions were obtained from the following resources: (a) literature curated resources such as the Human Transcriptional Regulation Interactions database (HTRIdb) (Bovolenta et al., 2012), Regulatory Network Repository (RegNetwork)(Liu et al., 2015), Transcriptional Regulatory Relationships Unravelled by Sentence-based Text-mining (TRRUST)(Han et al., 2015), TRANSFAC resource from Harmonizome (Rouillard et al., 2016), (b) ChEA3 containing ChIP-seq determined interactions(Keenan et al., 2019) and (c) high confidence protein-protein binding interactions (RF-RF) were retrieved from human protein-protein interaction network −2 (hPPiN2)(Ravichandran et al., 2021). The resulting interactions from both categories were collated and made a non-redundant set. This resulted in a connected set of RF-RF and RF-TG interactions that comprise a ‘master’ human Gene Regulatory Network (hGRN) with 27, 702 nodes and 890, 991 interactions **(Data file S1).** This extensive hGRN, which encompasses the experimentally determined interactions/edges, was used to infer stimulant-specific hGRNs and top paths using our inhouse network mining algorithm, ResponseNet. We have previously demonstrated that ResponseNet, which utilizes a knowledge-based network and a sensitive interrogation algorithm, outperformed data-driven network inference methods in capturing biologically relevant processes and genes. Furthermore, we have previously conducted a systematic comparison of our network mining strategy with other data-driven module detection methods, including jActiveModules (Ideker et al, 2002), WGCNA (Langfelder et al, 2008), and ARACNE (Margolin et al, 2006). Our findings demonstrated that our approach outperformed conventional data-driven network inference methods in capturing the biologically pertinent processes and genes (Ravichandran and Chandra, 2019).This was also validated by comparison with simulated data (Sambaturu et al., 2021).

### Model contexting and identification of epicentric RFset

Starting from the hGRN, stimulant-specific GRNs were derived by weighting the master hGRN network with differential transcriptomes of stimulants (ie., stimulant versus baseline) and generated weighted networks for individual stimulants. A sensitive network mining approach, previously established in the laboratory, was then applied to identify top-ranked activated and repressed sub-networks (ResponseNet) using Eq. 1, Eq. 2, and Eq. 3. This method of computing response networks which involves a knowledge-based network and a sensitive interrogation algorithm has been shown to outperform data-driven network inference methods in capturing biologically relevant processes and genes (Ravichandran and Chandra, 2019; Sambaturu et al., 2021). Briefly, the network mining algorithm works by computing minimum weight shortest paths, in which each path begins from a source node and ends with a sink node, identifying connected sets of edges that make up the least-cost paths. The shortest paths between all pairs of genes were computed using Dijkstra’s algorithm using Zen library implemented in Python2.7. For a path of length n, the path cost was calculated as a summation of the edge weights EW_ij_ of all edges forming the path, and normalized by the path length. Subsequently, all shortest paths were sorted based on their path costs from lowest to highest, and the top 1% were considered to constitute the top-perturbed network (TPN). Nodes forming the TPN were evaluated for enrichment of DEGs using Fisher’s exact test. Further, the epicentric nodes in each TPN were identified using the EpiTracer algorithm developed previously in the laboratory (Sambaturu et al., 2016). Epitracer estimates the influential state of a node by computing ripple centrality, which is the product of closeness centrality and outward reachability.

The TPN has two components, a top-activated network and a top-repressed network, generated by using appropriate node-weighting schemes. To generate the activated network, the weight of node i in a condition treated with Stimulant A (S_A_) was computed as:

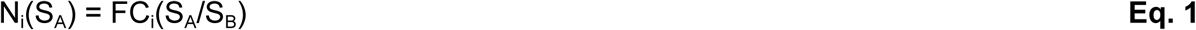

Where FC was the fold change of gene i in a condition treated with stimulant A (S_A_) with respect to the baseline S_B_ (antilog values were used to compute fold changes). To generate the repressed network, the node weight of node i in SA was computed as:

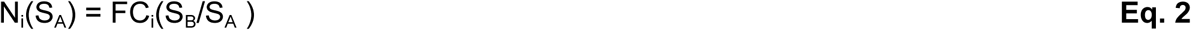

The edge weight EW_ij_ in a given condition S_A_ for an edge e comprised of nodes N_i_(S_A_) and N_j_(SA) was calculated as

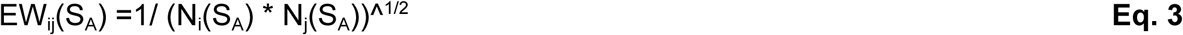

Where N_i_(S_A_) and N_j_(S_A_) are the node weights of nodes i and j, respectively. Lower the edge weight, stronger is the interaction between the nodes.

These subnetworks were combined to obtain a top perturbed network (TPN) for each stimulant. Nodes in the TPN were rank-ordered based on the ripple centrality score. Differentially expressed RFs with the ripple centrality score >0 were considered as influential transcription factors for a given TPN. The resultant RF-set defines the regulatory control for a given stimulant **(Fig. S13).**

### Clustering macrophage polarization states

We performed Monte Carlo reference-based consensus clustering using the M3C package implemented in R (John et al., 2020) to identify the optimal number of macrophage polarization states. Spectral clustering with Pearson’s correlation coefficient as a correlation metric for 200 iterations with 250 replicates was used to estimate the cluster structure that best fits the differential transcriptome of (a) entire transcriptome (n=12,164), and (b) RF-set1 (n=265). Further, Relative Cluster Stability Index (RCSI) and Monte Carlo adjusted p-values were computed using M3C to test the number and structure of the clusters. For the cluster visualization and comparison, Dendextend package in R (Galili, 2015) was used.

### Network visualization and Enrichment analysis

We have visualized networks using Cytoscape 3.2.0 in Allegro Spring-Electric layout (Shannon et al., 2003). We performed enrichment analysis for the genes of the regulatory cores of each cluster using Reactome pathway resource with default parameters(Fabregat et al., 2017) and the resultant hits with q-value ≤ 0.05 were considered to be significant (using a hypergeometric test).

### Reagents, Cell line and Bacteria

General laboratory chemicals were procured from Merck, Promega, Invitrogen, Himedia and Sigma-Aldrich. Tissue culture plastic wares were purchased from Biofil, BD Falcon and Tarson. Phorbol 12-myristate 13-acetate (PMA #P8139) and lipopolysaccharide (LPS #8630) were procured from Sigma-Aldrich. IL4 (#204-IL) were purchased from R&D Systems. THP-1 human monocytic cell line was obtained from National Centre for Cell Sciences, Pune, India and cultured in RPMI (GIBCO, Invitrogen Corporation, USA) supplemented with 10% heat inactivated FBS (Sigma-Aldrich, USA). THP-1 cells were treated with PMA (20ng/ml) for 24hrs for its differentiation into macrophages. Post PMA treatment, THP-1 derived macrophages were washed and kept in complete RPMI media followed by treatment/ infection as mentioned. *S. aureus Cowan 1* (MTCC - 902) was procured from Microbial Type Culture Collection, IMTECH Chandigarh. *S. aureus* was grown in Luria Broth at 37°C overnight. *S. aureus* was harvested and washed with phosphate-buffered saline (PBS) once the OD600 reached 0.6, and was then resuspended in the cell culture RPMI complete medium without antibiotics. *S. aureus* was used at 10 MOI for in vitro experiments.

### RNA isolation and quantitative real-time PCR

Total RNA was isolated from PMA differentiated THP-1 cells using TRI Reagent (Sigma-Aldrich, USA) as per the manufacturer’s protocol. For RT-PCR, 1μg of total RNA was converted into cDNA using First strand cDNA synthesis kit (BIO-RAD, USA) according to the manufacturer’s protocol. Quantitative real time PCR was performed with SYBR Green PCR mix (KAPABIOSYSTEMS, USA) for quantification of target gene expression. Amplification of housekeeping gene GAPDH was used as an internal control. The list of primers used for the experiment is provided in Table S2.

### Transient transfection

PMA differentiated THP-1 cells were transiently transfected with 100nM targeted siRNA using low molecular weight polyethyleneimine (Sigma-Aldrich, 40872-7). Further, 36 hrs post transfection, cells were treated or infected as indicated for required time and processed for analysis.

### *In vitro* CFU analysis

PMA differentiated THP-1 cells were infected with *S. aureus Cowan I* at an MOI of 10 for 2 hrs to facilitate internalization. Following 2 hours, cells were washed with complete DMEM (without antibiotics) and Gentamicin (100μg/ml) treatment was given for 2 hours to remove extracellular bacilli. Cells were lysed in 0.1% TritonX-100 containing 1X PBS and CFU estimation was done at 0hr (after gentamicin treatment) and 24hrs by plating appropriate dilution on LB plates.

### Statistical analysis

Levels of significance for comparison between samples were determined by the Student’s t-test distribution and one-way ANOVA followed by Tukey’s multiple-comparisons. The data in the graphs are expressed as the mean ± standard error for the values from at least 3 or more independent experiments and p values ≤ 0.05 were defined as significant. R version 3.6.3 was used for all statistical analyses.

## Results

### Subnetworks from contextualised hGRN identifies regulatory barcodes for macrophage polarization states

We first identified the ‘programmable space’ in macrophages in terms of the nature and range of transcriptional reprogramming that they achieve in response to different immunostimulants. The stimulants belong to broad classes of pro and anti-inflammatory cytokines such as Toll-like receptors (TLRs) ligands, immune complexes, interleukins, glucocorticoids or their selected combinations, comprehensively covering the macrophage polarization spectrum. We probed the programmable space by constructing and analyzing 28 unbiased ‘response networks’ of macrophages, corresponding to the treatment with 28 different immunostimulants. Towards this, we first constructed a knowledge-based human gene regulatory network (hGRN). Our hGRN covers regulatory elements from the whole genome and contains 27,702 genes and 890, 991 RF-RF and RF-TG directed interactions, which was rendered context-specific for each stimulant using their respective differential transcriptome data. We then subjected the 28 contextualized hGRNs to our network mining algorithm, which identified a connected set of genes that collectively vary the most in their expression levels in response to the given stimulant. This yielded 28 different subnetworks containing the highest ranked regulatory perturbations, with nodes ranging from 4032 to 6785 nodes **(Data file S2).** The resultant subnetworks containing connected sets of weighted and directed interactions capture cascades of regulatory events which are in essence the top-perturbed networks (TPNs) in each condition. Next, we configured a pipeline to identify the most influential transcription factors from the resultant TPNs. We used EpiTracer, an algorithm previously developed in the laboratory (Sambaturu et al., 2016) which identifies the highest ranked influential nodes in a given network using a ripple centrality measure. This step results in the identification of the genes that have the strongest ‘causal’ association with a given perturbation pattern. The word ‘causal’ here refers to the potential of the given gene for wielding an influence on the system, as estimated by the network analysis. Put together, we identified epicentric transcription factors (Ripple centrality score >0) which are differentially expressed in each of these TPNs. When pooled, these RFs from all stimulants result in a set of 265 RFs (**RF-set1 - Data file S3)**, defining the programmable space in relation to macrophage polarization.

Further, to probe the number of distinct polarization states, we carried out agnostic clustering based on the differential transcriptome of RF-set1, using a Monte Carlo reference-based consensus clustering (M3C) method. This resulted in thirteen different clusters. The clusters and the associated members are shown in **(Table S3).** The relative cluster stability index (RCSI) and Monte Carlo p-values further validated that the cluster stability **(Fig. S1A, B).** Put together, it is clear that the TPNs group into 13 broad clusters (**Fig. 1A**), each representing a distinct polarization state across the spectrum. Further, the cluster member associations inferred from RF-set1 were found to be concordant with the clustering pattern (**Fig. S2**, cophenetic coefficient = 0.68; p-value = 1.25e−51) inferred by considering the entire differential transcriptome (n=12,164). This clearly indicates that the RFs in RF-Set-1 are sufficient to characterize a given polarization state and that the subnetworks from the contextualized hGRNs define the macrophage polarization spectrum.

**Fig.1.**
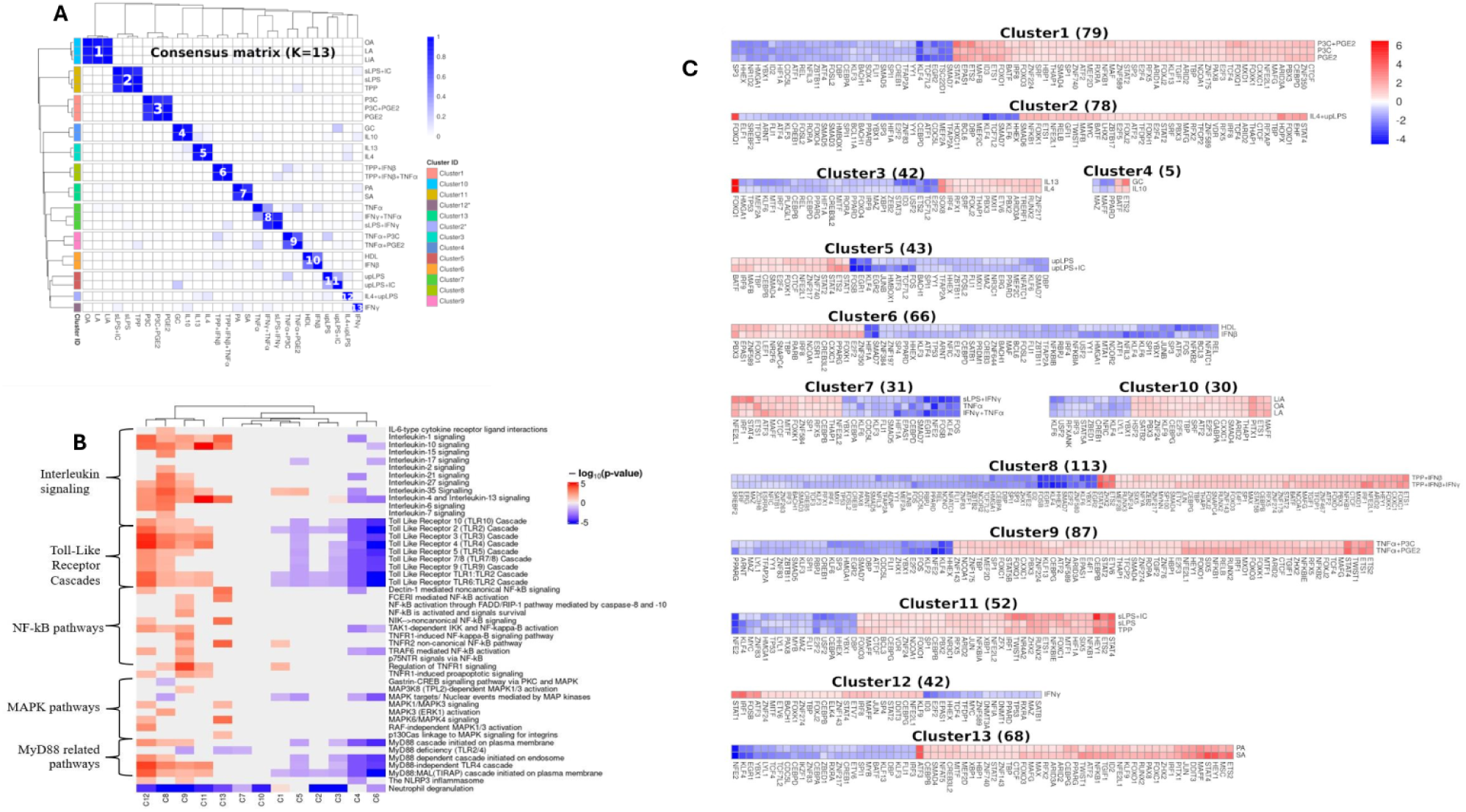
A clustergram showing K=13 for samples spanning the macrophage polarization spectrum and RF-barcodes characteristic of each polarization state. The clustering pattern was obtained by using M3C, a widely used agnostic clustering method based on Monte Carlo referencing that uses stability selection to estimate the number of clusters. (A) Cluster-member associations in the consensus matrix for K=13. Clusters with a single member are starred. Clusters 2 and 12 are singletons. (B) A heatmap showing pathways commonly enriched in inflammation and other immunological processes. The pathways marked in red are upregulated, while those marked in blue are downregulated. The metric chosen was the negative logarithm of the p-value for each associated pathway. (C) RF barcodes which are characteristic of the thirteen different clusters. Each barcode is a heatmap of the differential expression profile of the identified transcription factors (as compared to the unstimulated Mo state) in each cluster. The length of the barcode (which is the number of RFs in each cluster) is indicated within parentheses. Barcodes of members within a cluster resemble each other extensively and are shown together.

Functional enrichment analysis of the genes using pathways related to inflammation and other immunological processes from the Reactome pathway resource with default parameters (Fabregat et al., 2017) encompassing the regulatory cores of each cluster established their biological relevance. Clusters C11, C12, C8, C9 show strong activation of Interleukin signaling, Toll-Like Receptor cascades, as well as NF-kB pathways indicating a robust pro-inflammatory response (**Fig 1B**). In contrast, Clusters C3, C4 and C6 display significant downregulation of Toll-like receptor cascades and absence of interleukin signaling, suggesting suppression of the innate immune response. Next, we set out to identify combinations of key regulatory factors for each cluster. The subnetworks from which the RF-set1 and the subsequent clustering pattern was derived, serves as precise molecular definitions for each polarization state. Here, we seek to identify minimal gene sets from each cluster for identifying gene panels that are the most influential in guiding the differentiation into specific polarization destinations. Our ultimate goal was to probe if any of them could serve as switching factors between different functional states. The core networks of each cluster provide us connected sets of weighted and directed interactions which capture cascades of regulatory events. The complexity in these subnetworks arise due to the plurality of interactions between RFs since (i) a given RF can regulate multiple targets, (ii) multiple RFs can regulate a given target, and (iii) RFs themselves are regulated by other RFs. A computational pipeline was configured to discover combinations of transcription factors with specific expression patterns, from the subnetworks for each polarization state. The pipeline consists of computational steps to identify (a) common cores for each polarization state, and (b) differentially expressed epicentres of different members of the clusters. The first step was achieved by taking an edge-level intersection of the TPNs within each cluster, which resulted in pruning the subnetworks to retain only those nodes and edges that are common and exhibit the same pattern in all members within each cluster. The pipeline yielded a 52 RF-set characteristic of C11 and a 42 -RF set characteristic of C3 and similarly for all other states. These RF-sets serve as barcodes for each state where color gradient reflects the gene expression changes of each RF (as compared to an unstimulated state) (**Fig. 1C**). Put together, these barcodes amount to a union of 223 RFs (RF-set2) that can be considered to form a pool of central regulatory factors for the polarization states studied here (RF-set2 - **Data file S3**).

### Identification of a 3-RF set as M1 to M2 switching factors

We investigated if the barcodes could serve as switching factors to tune the polarization destination of macrophages. For this, we selected a stimulant that is known to drive unstimulated macrophages (Mo) to M1 and asked if we could switch it to an M2 destination instead. We start with the barcodes for both states (C11 - representing M1 and C3 representing M2) and pruned them to retain (a) only those genes that occur in the barcodes for both the states and (b) exhibit reversal in the gene expression variation trend between the two states. This resulted in identifying a panel consisting of sixteen RFs (*STAT4*, *ETS2*, *NFKB1*, *BCL3*, *NFE2L2*, *MTF1*, *HIF1A*, *NFKBIA*, *HEY1*, *FOXO3*, *ZFX*, *ZNF24*, *XBP1*, *CEBPB*, *MYC* and *NFE2)* where each gene exhibits a significant quantitative difference in gene expression variation between M1 and M2 states, both with reference to the Mo state (**Fig. 2C**). Of these, we selected the top-ranked genes based on the extent of gene expression variation in the two states and the number of targets they influenced. We identified two well-known RFs: *STAT4*, *NFKB1* and three additional RFs *NFE2L2*, *CEBPB*, *BCL3*, that qualified our criteria to be potential switching factors. STAT4 and *NFKB1* are well studied and clearly linked to the proinflammatory M1 state and have many targets. We verified that in our study system, they are indeed upregulated upon LPS induction and downregulated upon IL4 treatment (both with reference to Mo, **Fig. 2C**). We focused on the three newly identified RFs *NFE2L2, CEBPB, BCL3* (referred to as NCB hereafter), which show the same expression trend and are positioned to have a high degree of influence in the network. Put together, the 3-RF combination has 1293 targets, covering about 53% of the top-perturbed network. The parts of the respective top-perturbed networks containing the regulatory cores for C11 (M1) and C3 (M2) along with the associated pathways are shown in **Fig.2A** & **Fig.2B** respectively.

**Fig.2.**
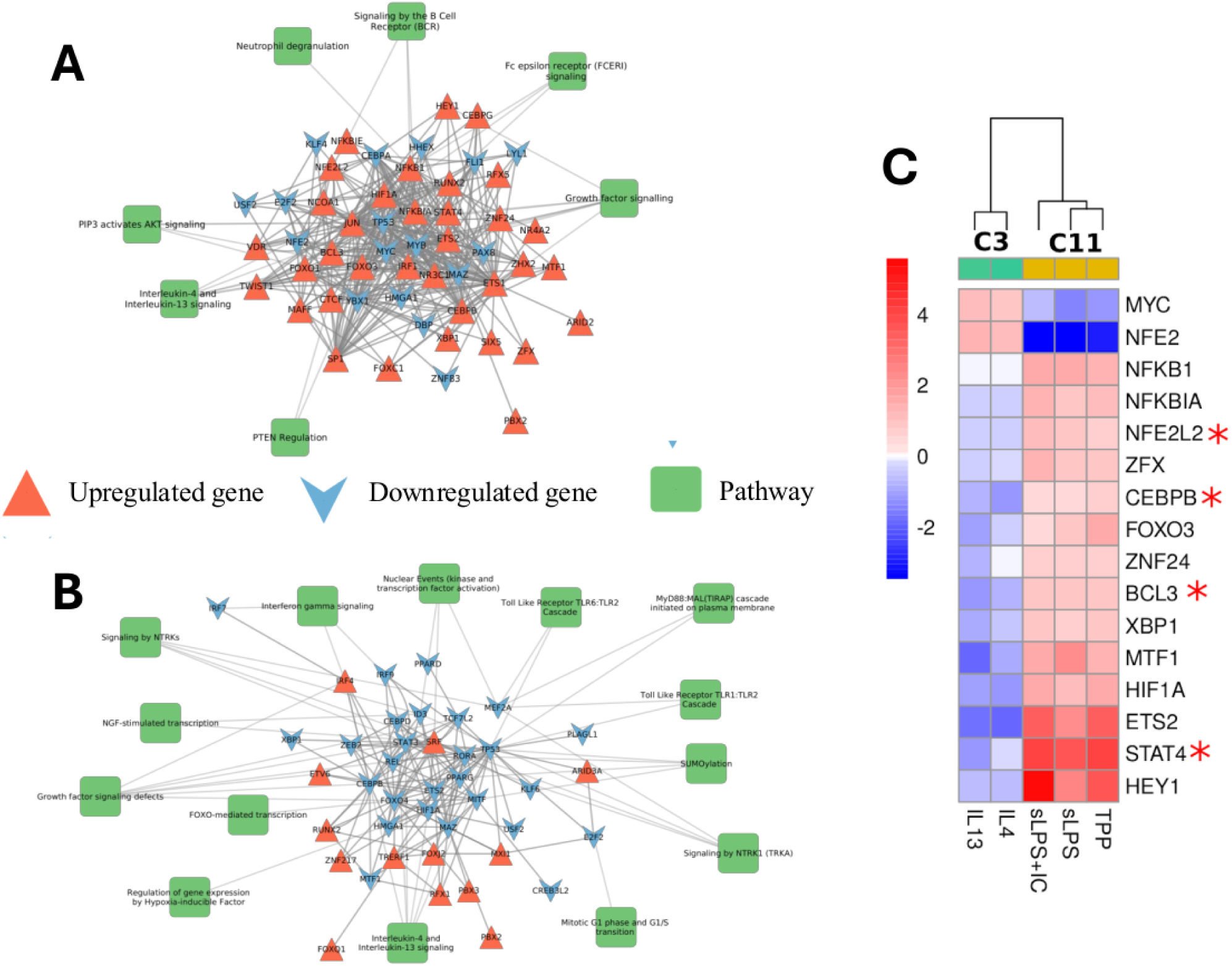
Switching factors that guide M1 to M2 destination. (A) The regulatory core for the M1 cluster (C11) showing the barcode genes. (B) The regulatory core for the M2 cluster (C3). The core networks contain the top perturbed regulatory interactions common across C11 members (stimulants: TPP, sLPS and sLPS + IC) and C3 members (stimulants: IL4 and IL13) respectively. (C) A heatmap of the genes in the barcode that exhibits a reversal in the fold change pattern between M1 (C11) and M2 (C3). RFs that are selected for experimental validation are starred.

We hypothesized that knock-down of *NFE2L2*, *CEBPB*, *BCL3* in an M1 state would switch it to M2. To test this hypothesis, we designed an experiment (experimental design shown in Fig. 3A) where we take THP-1 derived macrophages (taken as Mo) and stimulate them with sLPS to obtain an M1 state. We first verified that sLPS treatment is leading to an M1 state by measuring known M1 markers (*IL1B* in Fig. 3B), which confirmed that they were indeed in an M1 state. We also measured the levels of M2 markers (*IL10* in Fig.3B) and verified that there was no significant change. *STAT4* is known to have an effect on the polarization state as its knockdown in M1 was shown to significantly attenuate the M1 state (Ishii et al., 2009) and we therefore used *STAT4* as a positive control. We checked whether *STAT4* was exhibiting the expected behaviour in our experimental setup and we found that was indeed the case **(Fig. S3, Fig. 3C**). We then performed siRNA mediated combined knock-down of NCB, the 3-RF set, to evaluate the M1 to M2 switch. We verified that the siRNA mediated knockdown (referred to as siNCB) had worked by testing the gene expression values of the 3 NCB genes and finding them to be significantly reduced **(Fig. S4, Fig. 3D**). Upon stimulation with sLPS in the siRNA treated cells, we observed that the known M1 markers were significantly reduced while the known M2 markers were significantly pronounced, clearly showing that the siRNA knockouts lead the cells to shift from M0 to M2 (**Fig. 3D**). The change was statistically significant when compared with the parent THP-1 cells without knockdown as well as with non-targeting siRNA controls. In summary, we observe that the knockdown of the 3-gene set leads to a clear shift of the polarization state away from M1 towards M2 (significant decrease in the expression of *CXCL2, IL1B, iNOS*) and significant increase in classical M2 marker (*ARG1*), despite the sLPS trigger. In addition, we conducted siRNA knockdown experiments on the regulatory factors *NFE2L2, CEBPB,* and *BCL3*, and observed a similar trend of decreased expression of M1 markers and increased expression of M2 markers **(Fig. S5).** Among the three RFs, CEBPB had the most significant effect on the switching of M1 and M2 marker expression, with significant decreases in the expression of *CXCL2* and IL1B (M1 markers) and significant increases in the expression of *ARG1, IL10,* and *MRC1* (M2 markers). In comparison to *STAT4* (positive control – Fig. 3C), knockdown of *CEBPB, NFE2L2*, and *BCL3* had significantly higher effects on the expression of M1 and M2 markers, indicating their role in modulating the inflammatory response.

**Fig. 3.**
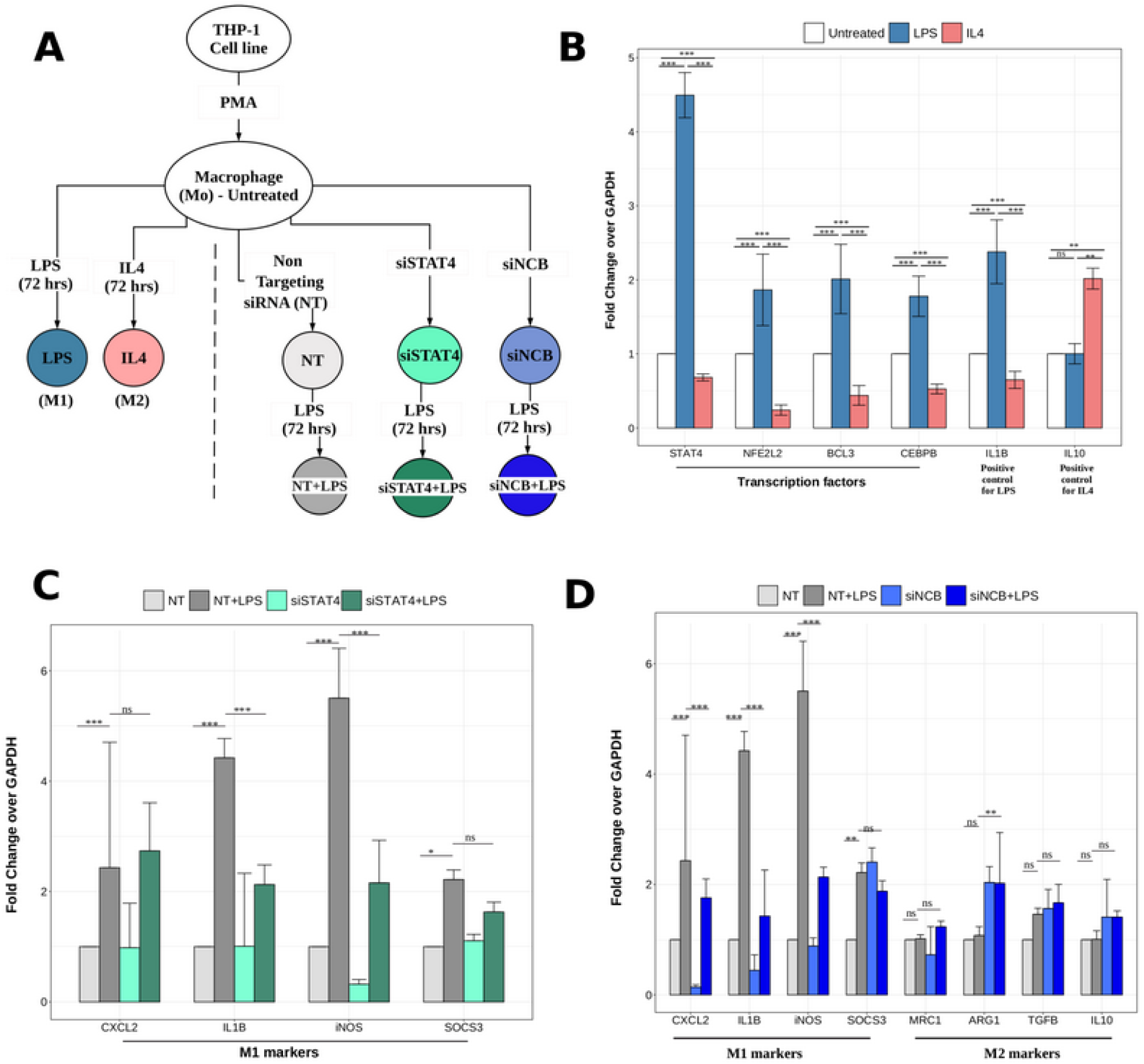
siRNA knockdown of *NFE2L2*, *CEBPB* and *BCL3* in LPS infected THP-1 derived macrophage cell line attenuates macrophages polarizing to C11 state (M1) (A) Schema describing the experimental design for testing the role of the 3-gene set NCB, predicted to be a switching factor combination between M1 and M2 destinations. The LPS and IL4 treatment conditions are included to verify the experimental setup. The non-targeting siRNA is used as a negative control while siSTAT4 is used as a positive control. siNCB is a knock-down of the 3-gene set (NCB) which is evaluated in this study. (B) Gene expression changes of selected RFs and known markers of M1 (IL1B) and M2 (IL10) in THP1 cell lines upon LPS and IL4 treatment (for 72 hr) respectively. Color coding scheme is similar to that in (A). THP1 monocytes were treated with PMA (20ng/ml) for 16h. Post 24h, LPS (100ng/ml) and IL-4(1000IU) treatment was given for 72 hr. Two-way ANOVA (P > 0.05 :ns, P < 0.05:*, P < 0.01:**, P < 0.001:***; N=3 biological replicates) is used for estimating significance in gene expression fold change normalized over untreated cells. (C) Gene expression changes in the M1 markers upon siRNA knockdown of STAT4 (positive control). THP1 monocytes were treated with PMA (20ng/ml) for 16h. STAT4 specific siRNA was transfected. Post 8h, LPS (100ng/ml) treatment was given for 72h. Gene expression changes in M1 markers were quantified (Two-way ANOVA; P > 0.05 :ns, P < 0.05:*, P < 0.01:**, P < 0.001:***; N=3 biological replicates; NT: non-targeting siRNA) (D) Gene expression changes in the M1 and M2 markers upon siRNA knockdown of the NCB set. THP1 monocytes were treated with PMA (20ng/ml) for 16h. NCB (*NFE2L2*, *CEBPB*, *BCL3*) specific siRNA was transfected. Post 8h, LPS (100ng/ml) treatment was given for 72h. Gene expression changes of M1 and M2 markers were quantified (Two-way ANOVA; P > 0.05 :ns, P < 0.05:*, P < 0.01:**, P < 0.001:***; N=3 biological replicates; NT: non-targeting siRNA)

In addition, we also identified a group of regulatory factors that may be involved in switching macrophages from an M2 functional state to an M1 state using the similar approach. These factors include *FOXQ1, SOX8, ETS2, MTF1, HIF1A, CEBPB, XBP1, IRF7*, and *TCF7L2* **(Fig. S6)**.

### Knockdown of NCB restricts the growth of *S. aureus* within macrophages

*S. aureus* is a facultative intracellular pathogen causing a broad spectrum of diseases such as minor skin infections to serious bloodstream infections and endocarditis. This pathogen is known to survive inside the host cell and escapes from professional phagocytic cells such as macrophages through various immune evasion strategies, especially through cytoprotective mechanisms by altering the host gene expression (Koziel et al., 2009)(Pidwill et al., 2020).

We performed differential analysis of peripheral macrophages infected with *S. aureus* using the publicly available data (Koziel et al., 2009), and profiled the expression levels of RFs in M1 (C11) barcode and M1 markers. This analysis showed an upregulation of NCB and other RFs in the M1 barcode at the 8hr, 24 hr and 48hr time point **(Fig. S8).** We then tested the same in an *in vitro S.aureus* infection model using THP-1 macrophages and made a very similar observation, where the mRNA expression levels of the STAT1, NCB, M1, and M2 markers were quantified at 12hr, 24hr, 48hr and 72hr time points **(Fig S10, S11).** Specifically, the expression levels of *NFE2L2, CEBPB, and BCL3* were significantly upregulated in the THP-1 macrophages upon *S. aureus* infection with respect to the uninfected control at 24 hrs **(Fig. S11).** This suggests that macrophages infected with *S.aureus* polarize to the M1 state within 24 hrs. Hence, this time point was used for further experiments.

Next, to evaluate the potential of NCB in regulating polarization of macrophages during *S. aureus* infection, we used the same in vitro infection setup but using the NCB knockdown cells (We ascertained that these RFs were sufficiently responsive at the selected time point of 24 hrs **(Fig. S9 A,B,C)).** We observed a significant shift from M1 to M2 with the downregulation in the expression of M1 markers (*CXCL2, IL1B, iNOS, SOCS3*) (**Fig. 4B, Fig. S12)** and upregulation in the expression of M2 markers (*TGFB* and *IL10*) (**Fig. 4B**). Interestingly, we observed 1.6 fold significant increase in the burden of S.aureus within these macrophages (**Fig. 4C**). This clearly indicates that knockdown of NCB combination renders the macrophage more conducive for *S. aureus* to proliferate.

**Fig. 4.**
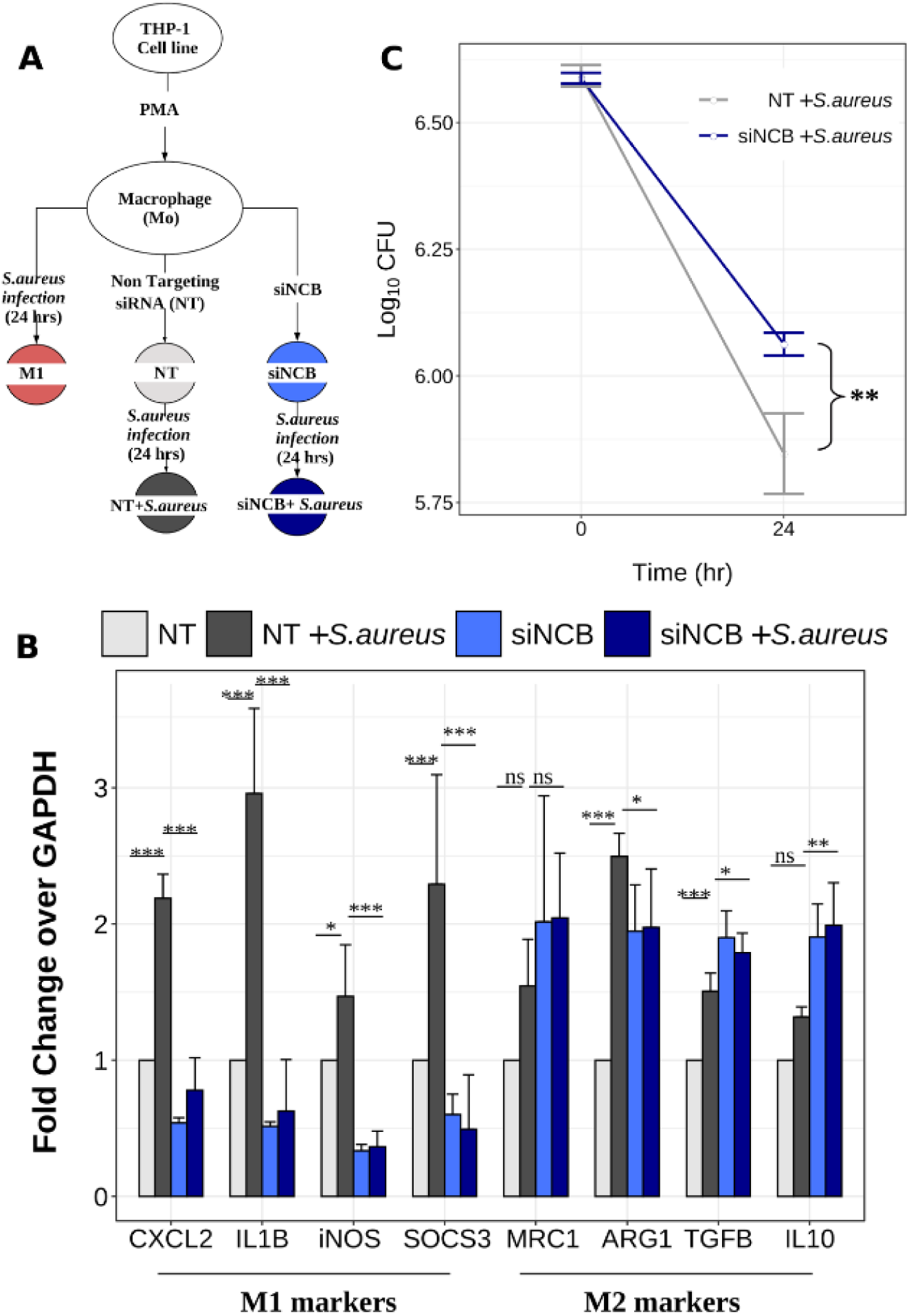
siRNA knockdown of NCB (NFE2L2, CEBPB and BCL3) in S.aureus infected macrophage increases infectivity. (A) Schema describing the experimental design. (B) Effect of the NCB set knockdown in the S. aureus infection model. siRNA knockdown of NFE2L2, CEBPB and BCL3 in THP-1 cells infected with S. aureus leads to a significant decrease in the gene expression of the M1 markers (*CXCL2, IL1B, iNOS and SOCS3*) and an increase in the expression of the M2 markers (TGFβ and IL10). Color coding is based on the schema shown in (A). THP1 monocytes were treated with PMA (20ng/ml) for 16hrs. NFE2L2, CEBPB and BCL3 specific siRNAs were transfected. Post 24hrs, S.aureus infection at 1:10 MOI was given for 24hrs. Gene expression levels of M1 and M2 markers were assessed. (Two-way ANOVA; P > 0.05 :ns, P < 0.05:*, P < 0.01:**, P < 0.001:***; N=3 biological replicates; NT: non-targeting siRNA). (C) In vitro S.aureus CFU assay. THP-1 cells were differentiated to macrophages with PMA for 16 hrs. Post 24 hrs, siRNA specific for NFE2L2, BCL3 and CEBPB were transfected. S.aureus infection (MOI 10) was given for 2 hrs prior to 2 hrs gentamicin treatment. CFU assay was performed at 0 hr and 24 hrs. (Two-way ANOVA; P > 0.05 :ns, P < 0.05:*, P < 0.01:**, P < 0.001:***; N=3 biological replicates; NCB : *NFE2L2, CEBPB and BCL3*). CFU-Colony Forming Units.

## Discussion

Macrophages display a wide array of functions, from being killers of pathogens to healers of damaged tissue (Murray, 2017; Murray and Wynn, 2011). Their versatility is attributed to their ability to respond to specific physiological or pathological cues, leading to their polarization into different states, each with a distinct functional phenotype. A range of molecular triggers that cause differentiation in macrophages is known (Hu et al., 2021; Jaguin et al., 2013; Xue et al., 2014). It is also well known that the differentiation spans a full spectrum of phenotypes with the classically activated M1 and the alternatively activated M2 states at two ends of the spectrum. The molecular profiles that define or cause these states, however, have not been sufficiently characterized. In particular, very little is known about the regulatory genes that control and guide the polarization in a specific direction. It is of great interest to identify RFs that are most influential in guiding the polarization. Given the importance of macrophages, several studies have investigated the variations in the transcriptomes in different diseases (Benoit et al., 2008; Mosser and Edwards, 2008; Ross et al., 2021; Schultze, 2016; Zheng et al., 2018) as well as upon exposure of macrophages to different exogenous and endogenous stimuli (Hu et al., 2021; Jaguin et al., 2013; Xue et al., 2014), which have resulted in a wealth of data that are publicly accessible. These large datasets provide opportunities for addressing a variety of questions.

In this study, we algorithmically identify distinct polarization states, minimal regulatory signatures for each state as transcription factor barcodes and demonstrate that manipulation of the barcode leads to loss of switching, in an example case. Additionally, our work has indicated that arresting the switch to M1 leads to hypersusceptibility to *S.aureus* infection in cell lines, suggesting that M1 has a distinct role in limiting the extent of infection. Two aspects in our computational workflow are distinct from the previous methods, which facilitated the identification of barcodes characteristic of different macrophage polarization states. The first is the use of an unbiased knowledge-based gene regulatory network as compared to the more common correlation-based interaction networks. In the latter, the interactions are statistical associations, with no guarantee that the interaction is ‘real’ in the given biological system (Petri et al., 2015; Ravichandran and Chandra, 2019). We overcome this limitation by consolidating a comprehensively curated knowledge-based network where each interaction is based on experimental data reported in databases and in primary literature. We have previously carried out a systematic comparison and have shown that knowledge-based networks are superior to correlation-based networks in capturing biologically relevant genes and pathways (Ravichandran and Chandra, 2019). Our macrophage specific GRN (subset of hGRN) network contains 326, 281 interactions among 742 RFs and 11, 423 target genes, which is more comprehensive than the previously reported knowledge based macrophage GRNs in literature (Hörhold et al., 2020; Palma et al., 2020). The second is the network interrogation method. Our method of contextualizing the knowledge-based network for diverse stimulants using the corresponding transcriptomes allowed a quantitative comparison of variations over a common background network, yielding a top-perturbed regulatory subnetwork in each case. Our network interrogation method identifies the most influential ‘epicentric’ transcription factors, which lead to the derivation of quantitative barcodes of RF panels. To the best of our knowledge, such barcodes have not been reported earlier even for the well-studied M1 and M2 states. We are hopeful our results of epicentric RFs and their molecular interaction trajectories will serve as key inputs for several investigations on the RFs individually as well as for specific RF panels. The response networks for each stimulant also serves as a rich resource of the top-ranked regulatory pathways in each case.

As a first step into the identification of barcodes, we used a pooled set of differentially expressed epicentric RFs (n=265) in different conditions that we identify from the networks and perform an unbiased clustering of the macrophages. We used a rigorous clustering method, which produced a robust clustering pattern which clearly indicated that the 28 stimulants group into thirteen polarization states. A clustering pattern was described earlier for the same dataset based on the top 1000 most varying DEGs from each condition, which resulted in nine distinct states (Xue et al., 2014).The clustering pattern that we get here from the top-ranked epicentres in each, yielded a similar pattern, indicating that a subset of DEGs that occur in our topnets are sufficient to achieve the differentiation. In fact, our gene-regulatory models implied that the saturated (SA and PA) and unsaturated (LA, LiA, and, OA) fatty acids mediate different modes of resolution that were grouped together in the previous clustering pattern, and are now resolved into two sub-branches. Similarly, the effects of IFNγ and sLPS which were clubbed in the previous pattern are now resolved, consistent with the known regulatory differences (Hoeksema et al., 2015; Kang et al., 2019). The principal takeaway from this analysis is not the exact number of clusters but rather the molecular basis it provides for the differentiation of functional states, with M1 and M2 representing two ends of the spectrum. Several other states are dispersed within the polarization spectrum, which can be described as a punctuated continuum. For our switching studies, we focused on clusters C11 (M1-like) and C2 (M2-like) due to their established functional relevance. However, future studies are required to explore the functional relevance of other clusters.

Our results suggest that the barcodes could serve as the regulatory handles to attain different polarization states with different functional phenotypes. We test that hypothesis for the M1/M2 axis and indeed demonstrate that siRNA knockdown of the RFs in the barcode achieves polarization state switching from M1 to M2 polarization states. In the M1 barcode, we selected the RFs upregulated as compared to the baseline (M_0_): (NBC-Nuclear Factor, Erythroid 2 Like 2 (*NFE2L2*), B-Cell Lymphoma-3 (*BCL3*), and CCAAT/enhancer-binding protein beta (*CEBPB*)). The roles of NFE2L2 and BCL3 in inflammatory responses in general have been discussed earlier (Wessells et al, 2004; Kobayashi et al, 2016). By knocking down the expression of these transcription factors in LPS treated THP-1 derived macrophage cell line, the cells were clearly seen not only to have not attained an M1 state but to have shifted towards the M2 state. This establishes their role as a switch between the states-where an up-regulation moves them from an M_0_ to M1 while a knock-down shifts them from M_0_ to M2, even after exposure to a M1-stimulant, thus indicating them to be essential for establishing the M1 state. Individual knockdown of the three genes was also performed to assess the contribution of each gene. It was observed that knocking down *CEBPB* led to a significant downregulation in the expression of M1 markers and an increase in M2 marker expression. In contrast, NFE2L2 and BCL3 knockdown resulted in decreased expression of M1 markers without a corresponding significant increase in M2 markers. These results suggest that *CEBPB* is crucial for M1 to the M2 transition. After having established an arrest of polarization to M1, we were curious to see if it resulted in a functional phenotype. Our experiments with *S.aureus* infection, clearly showed that the NBC knockdown led to increased infectivity of *S.aureus*. Expression of these RFs and their target genes are involved in cellular response to stress, Tumor Necrosis Factor (TNF) signaling pathway and senescence associated secretory phenotype, suggesting that inhibition of inflammatory associated processes halts M1 phenotype and promotes the M2-like Phenotype, which renders them hyper-susceptible to *S.aureus* infection.

In conclusion, we report a systems level analysis of the gene regulatory networks to identify regulatory factors which define different macrophage activation states. Perturbation of these factors for reprogramming the macrophage subpopulations to desired polarization states can be explored for applications in controlling the inflammation gradient in different diseases such as sepsis, autoimmune conditions and chronic infections.

## Supporting information

All supplementary data

## Author contributions

SR and NC conceptualized the study. KNB planned the validation experiments. SR carried out the data curation, computational methodology, analysis, and interpretation. BB and AS carried out the experiments. DD contributed to the data analysis and manuscript revision. NC and KNB acquired funding for the research. NC supervised the whole project. SR and NC wrote the manuscript with inputs from KNB BB, and AS. All authors read and approved the final manuscript.

## Acknowledgements

This work was supported by funds from the Department of Biotechnology BIC and NNP projects (SP|DBT0-24-0023, SP|DBT0-20-0025) and Department of Biotechnology Department of Science and Technology - Fund for Improvement of S&T Infrastructure in Higher Educational Institutions (DST-FIST), the University Grants Commission, and the Department of Biotechnology (DBT), Government of India; (DBT)-Indian Institute of Science partnership program (Phase-II at IISc, BT/PR27952/INF/22/212/2018); (DBT, No. BT/PR13522/COE/34/27/2015 and BT/PR27352/BRB/10/1639/2017 to KNB) and the Department of Science and Technology (DST, EMR/2014/000875 to KNB). K.N.B. also thanks the Science and Engineering Research Board (SERB), DST, for the award of J. C. Bose National Fellowship (JBR/2021/000011 and SB/S2/JCB-025/2016) for the requisite funding support.

## Supplementary materials

**Fig S1.** M3C based identification of the optimal number of clusters for the entire transcriptome.

**Fig S2.** Clustering based on RF-set1 Vs. Entire transcriptome.

**Fig S3.** Validation of STAT4 siRNA.

**Fig S4.** Validation of NCB siRNA.

**Fig S5**. Validation of CEBPB, NFE2L2 and BCL3 siRNA

**Fig S6.** Switching factors that guide M2 to M1 destination

**Fig S7:** Principal Component Analysis results for the 13 clusters based on hallmark genes of inflammation (S7A) and the NCB gene panel alone (S7B)

**Fig S8.** S. aureus infected macrophages polarize to M1.

**Fig S9:** Expression change of NFE2L2 (S9A), CEBPB(S9B) and BCL3(S9C) across time for M1 and M2 associated stimulants in the existing dataset

**Fig S10.** Gene expression changes in M1 and M2 markers upon S.aureus infection at different time points.

**Fig S11.** Gene expression changes in STAT4 and NCB-set upon S.aureus infection at different time points.

**Fig S12.** NCB (NFE2L2, CEBPB, BCL3) siRNA validation in S.aureus data.

**Fig S13:** Schematic diagram explaining the ResponseNet and EpiTracer algorithms

## Supplementary tables

**Table S1.** In vitro stimulant data.

**Table S2.** The list of primers used for the qRT-PCT

**Table S3.** The list of clusters, associated members, network details of regulatory cores and the number of RFs in the barcodes.

## Supplementary data file

**Data file S1:** The human Gene Regulatory Network.

**Data file S2:** Twenty eight different subnetworks capturing highest ranked regulatory perturbations.

**Data file S3:** RF-set1, RF-set2

