## Supplementary material for "Knock-down of a regulatory barcode shifts macrophage polarization destination from M1 to M2 and increases pathogen burden upon *S. aureus* infection": All supplementary data

**Supplementary figures:**

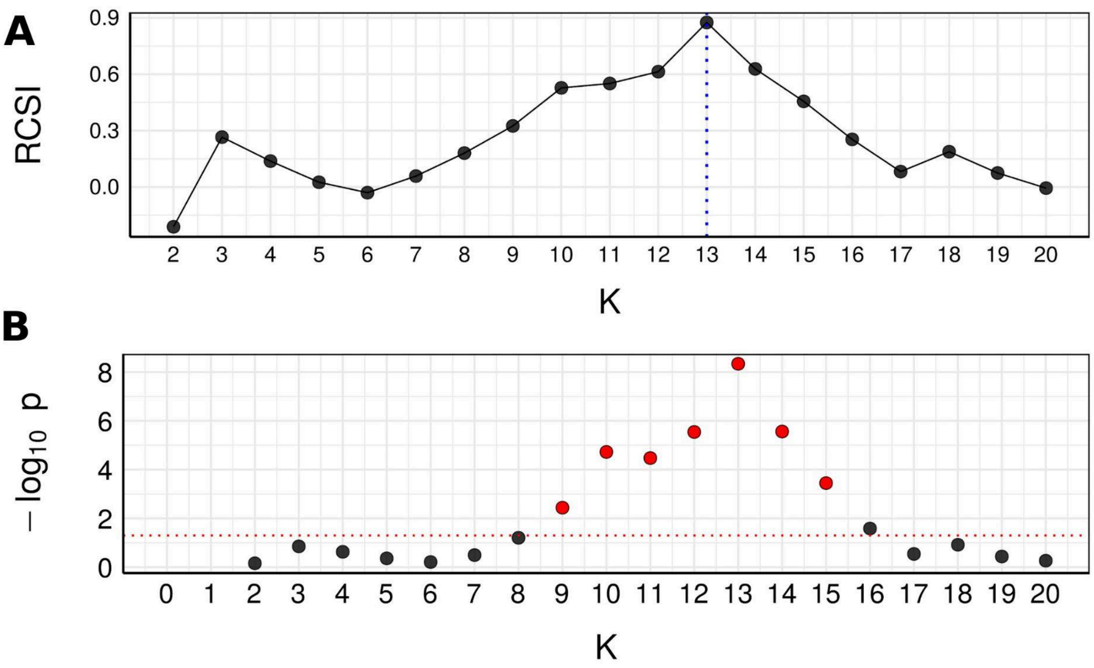


**Fig S1: M3C based identification of the optimal number of clusters for the entire transcriptome.**

Clustering pattern based on the differential profile of the entire transcriptome (n=12,164) showed thirteen different clusters spanning the macrophage polarization spectrum based on the (A) RCSI (spiked at K=13) with (B) significant p-value.

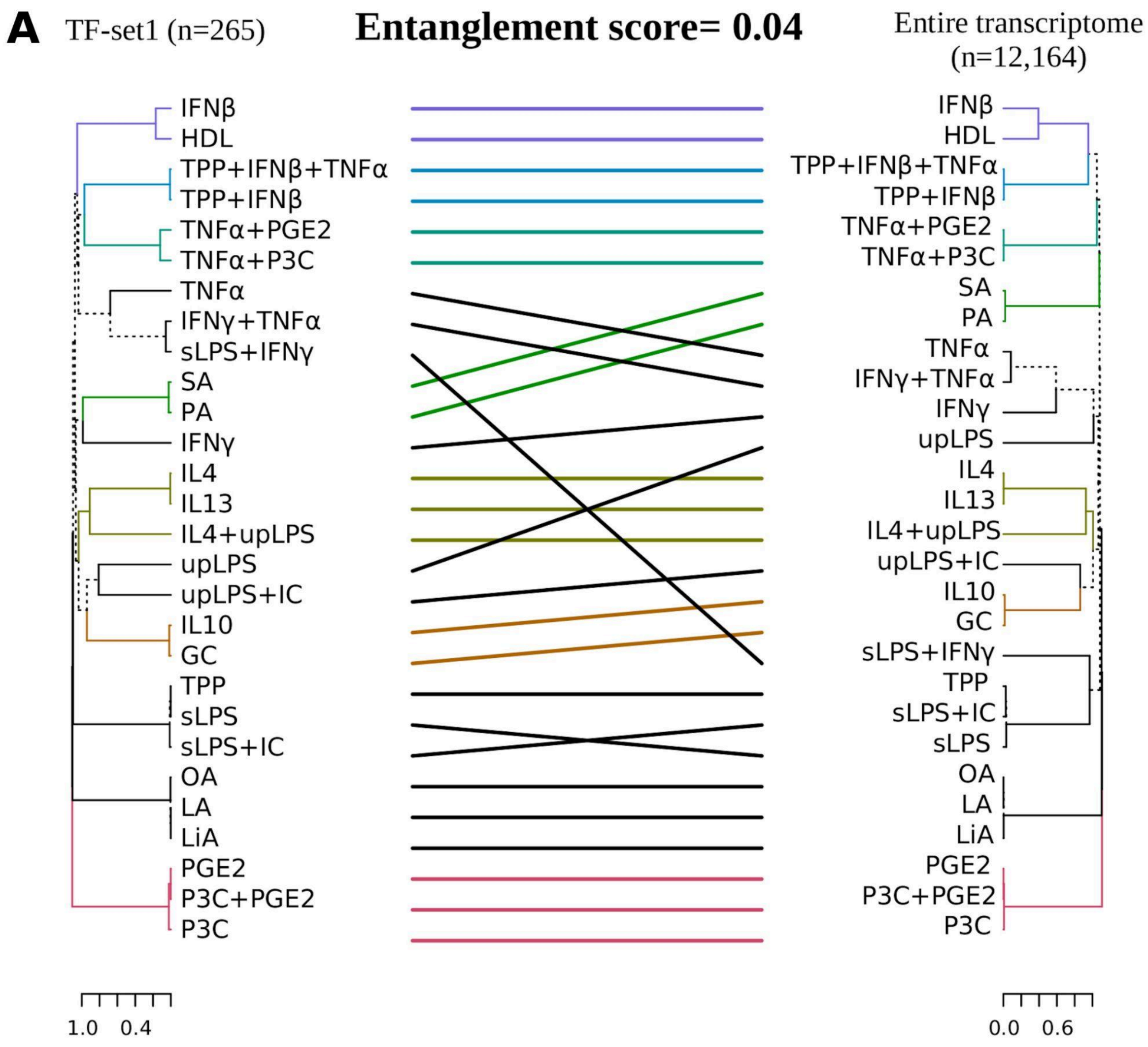


**Fig S2: Clustering based on TF-set1 Vs. Entire transcriptome.**

Entanglement score ranges from 0 to 1. Lower the score, better the alignment.

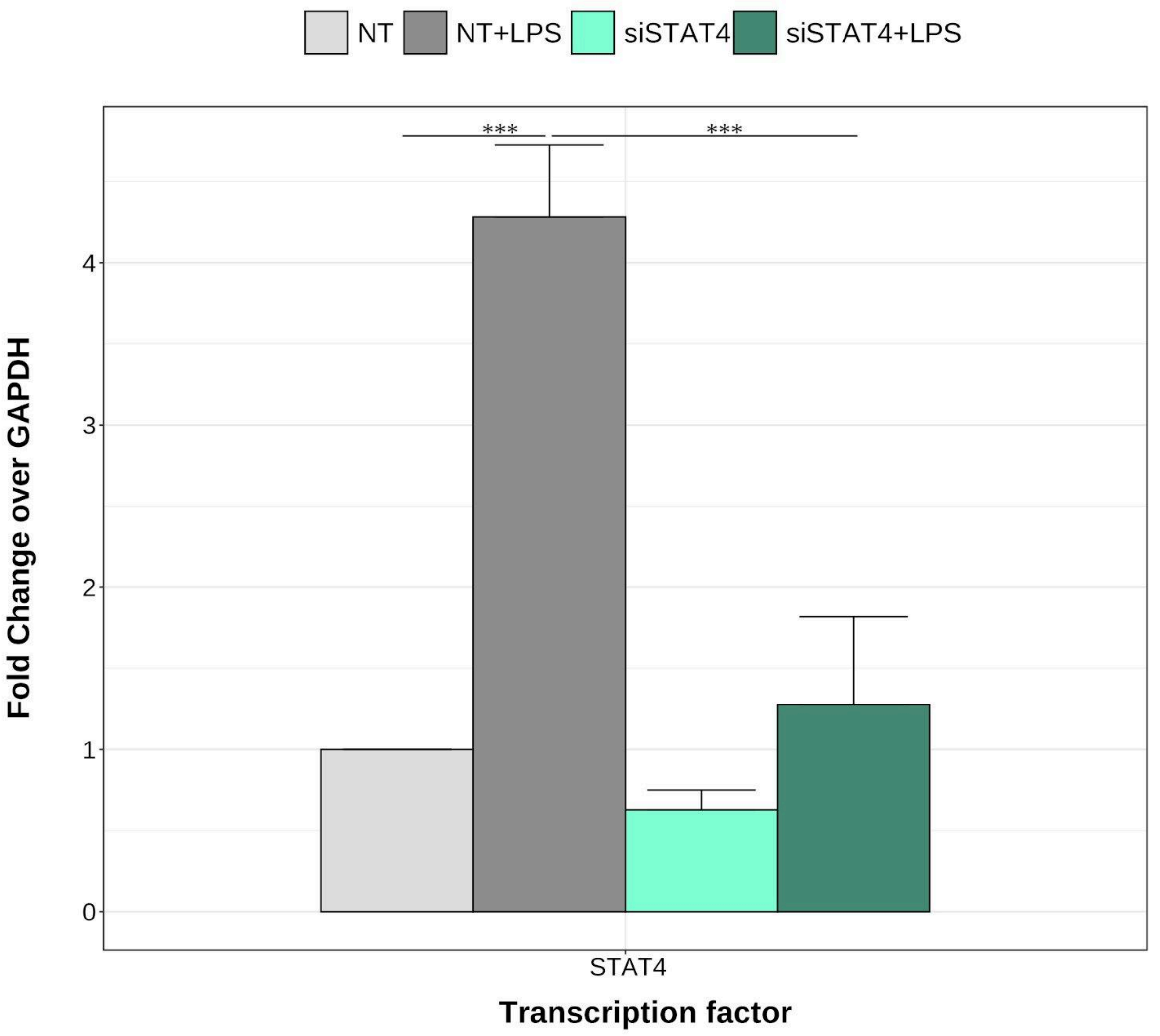


**Fig S3: Validation of STAT4 siRNA.**

THP1 monocytes were treated with PMA (20ng/ml) for 16h. STAT4 specific siRNA was transfected. Post 8hr, LPS (100ng/ml) treatment was given for 72hr. Transcript levels of STAT4 were assessed. {Two-way ANOVA,P > 0.05 :ns, P < 0.05 :*, P<0.01 :**, P<0.001 :***, N=3, NT: non-targeting siRNA}

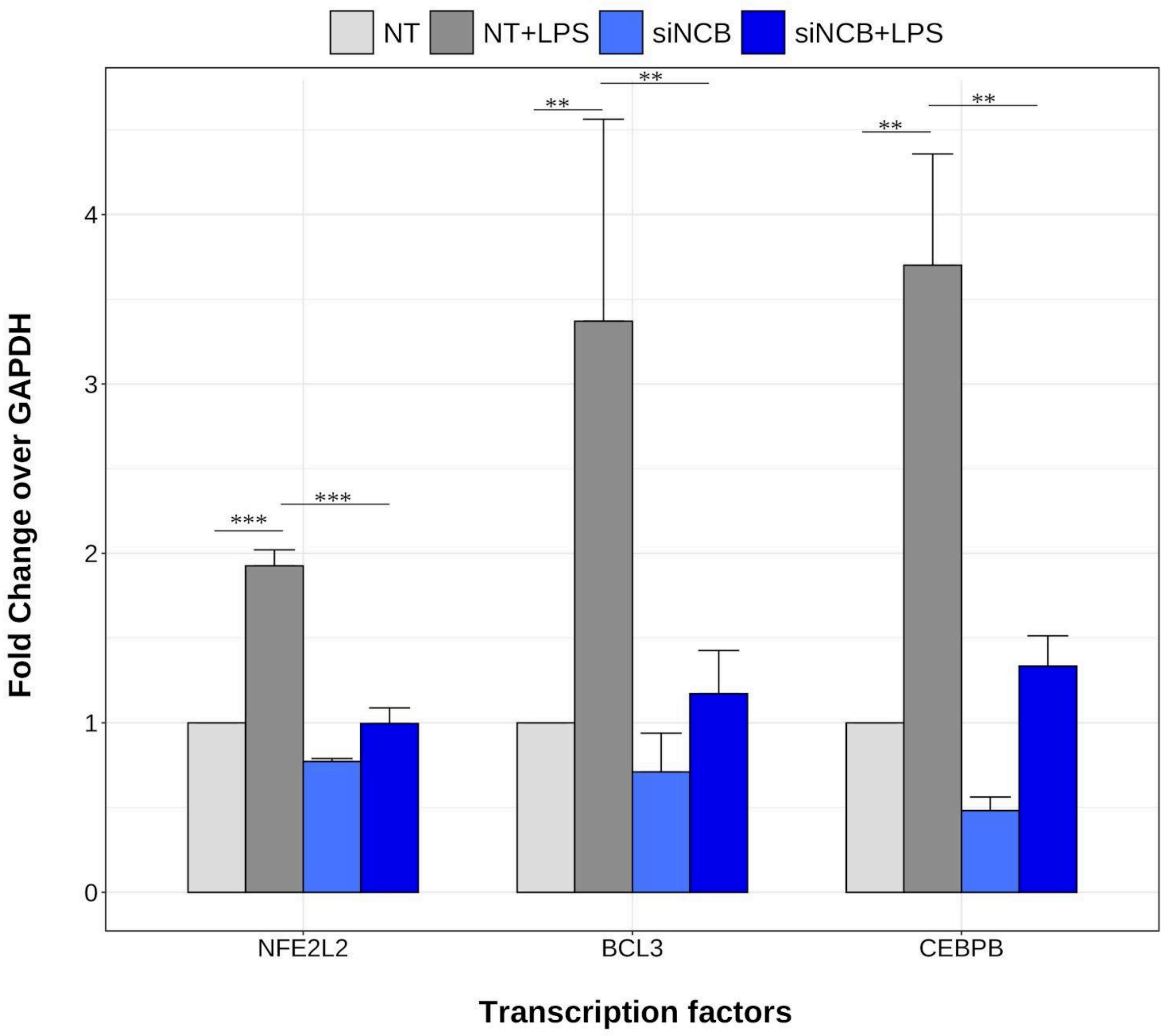


**Fig S4: Validation of NCB siRNA.**

THP1 monocytes were treated with PMA (20ng/ml) for 16h. NCB (NFE2L2, CEBPB , BCL3) specific siRNA was transfected. Post 8hr, LPS (100ng/ml) treatment was given for 72hr. Transcript levels of STAT4 were assessed. {Two-way ANOVA,P > 0.05 :ns, P < 0.05 :*, P<0.01 :**, P<0.001 :***, N=3 biological replicates, NT: non-targeting siRNA}

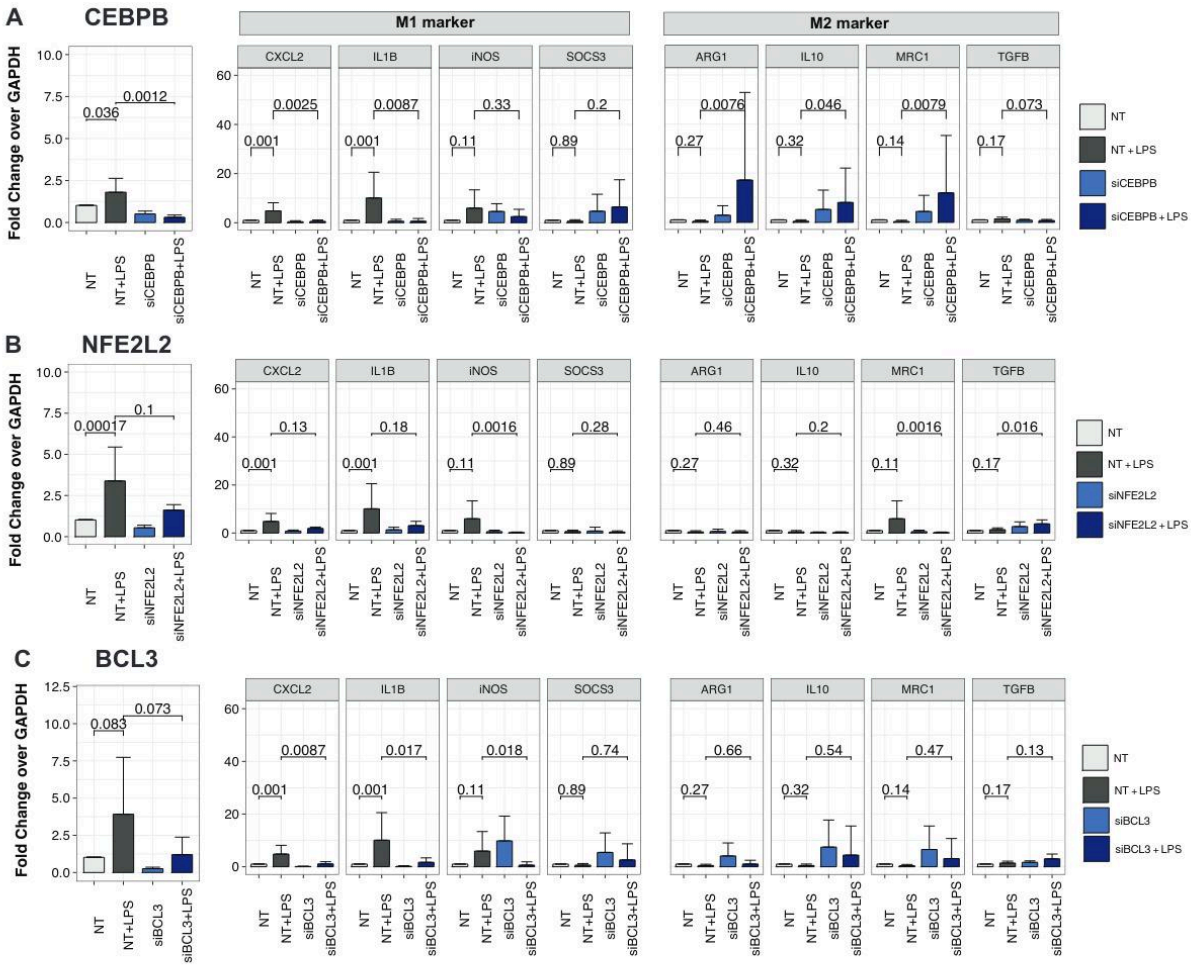


**Fig S5: Validation of CEBPB, NFE2L2 and BCL3 siRNA**

(A). Validation of CEBPB siRNA. Gene expression changes of selected known markers of M1 (CXCL2, IL1B, iNOS and SOCS3) and M2 (ARG1, IL10, MRC1 and TGFB) upon siRNA knockdown of CEBPB. Color coding scheme is similar to that in (Fig 2A). (B). Validation of NFE2L2 siRNA. Gene expression changes of known M1 and M2 markers upon siRNA knockdown of NFE2L2. (C) Validation of BCL3 siRNA. Gene expression changes of known M1 and M2 markers upon siRNA knockdown of BCL3. {Two-way ANOVA, $P > 0.05$ :ns, $P < 0.05$ :*, $P<0.01$ :**, $P<0.001$ :***, N=3 biological replicates, NT: non-targeting siRNA}

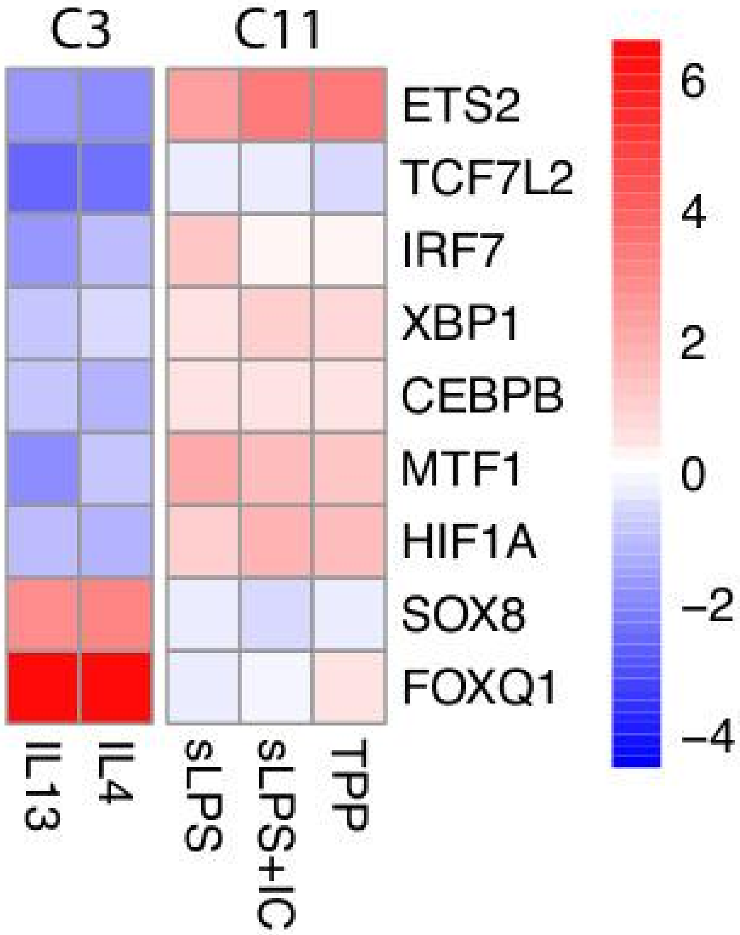


**Fig S6: Switching factors that guide M2 to M1 destination.**

A heatmap of the genes in the barcode that exhibits a reversal in the fold change pattern between M2 (C3) and M1 (C11)

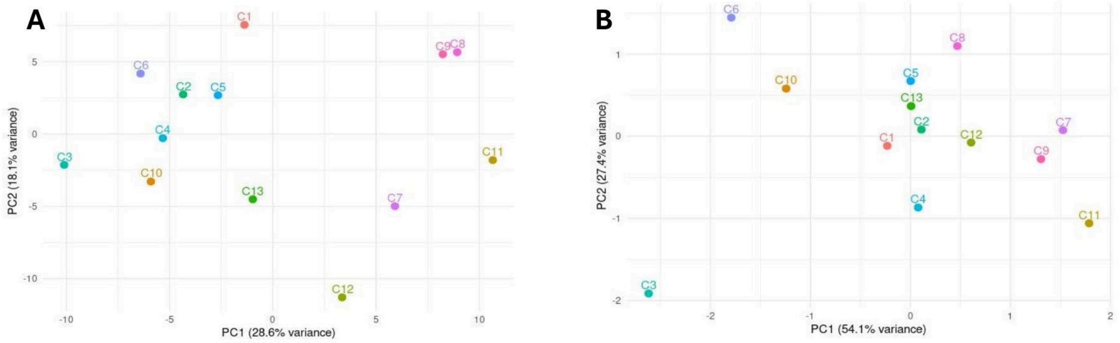


**A:** PCA plot segregating the 13 clusters based on the fold change values of hallmark genes of inflammation derived from MSigDB
**B:** PCA plot segregating the 13 clusters based on the fold change values of the NCB gene panel

**Fig S7: Principal Component Analysis of the thirteen clusters.**

(A) PCA plot segregating the 13 clusters based on the fold changes of hallmark genes of inflammation derived from MSigDB. (B) PCA plot segregating the 13 clusters based on the fold changes of the NCB gene panel.

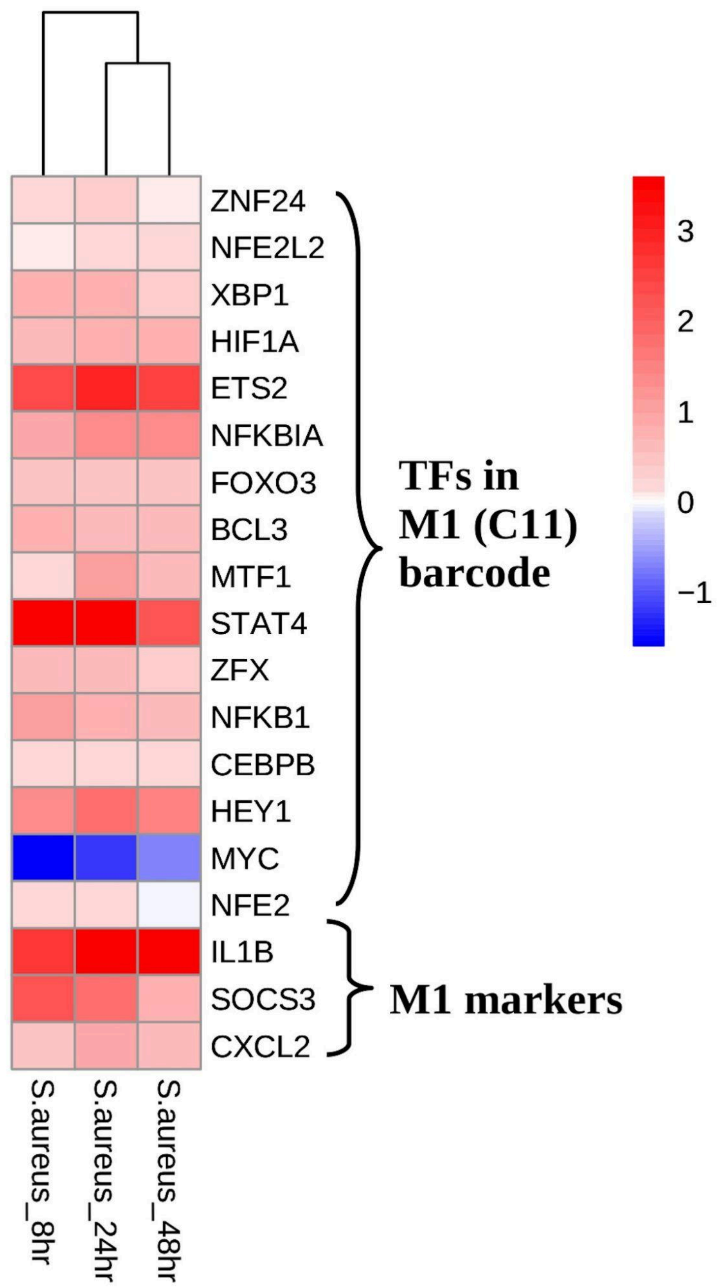


**Fig S8: S. aureus infected macrophages polarize to M1.**

Heatmap showing the differential transcriptome profile of 16 TFs in C11 barcode (M1) and M1 markers (IL1B , SOCS3, CXCL2) in S. aureus infected peripheral macrophages at 8hr, 24 hr and 48 hr.

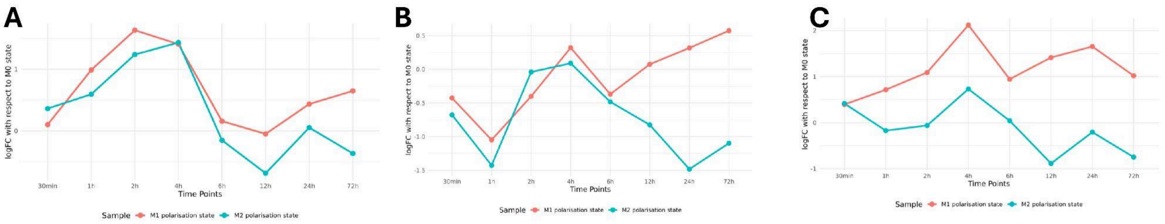


Gene expression changes at different points of **(A)** NFE2L2, **(B)** CEBPB and **(C)** BCL3 during stimulation of M0 cells with M1 and M2 associated stimulants

**Fig S9: Gene expression change of the NCB gene panel during stimulation of M0 cells with M1 and M2 associated stimulants respectively.**

(A) Gene expression change of NFE2L2. (B) Gene expression change of CEBPB. (C) Gene expression change of BCL3.

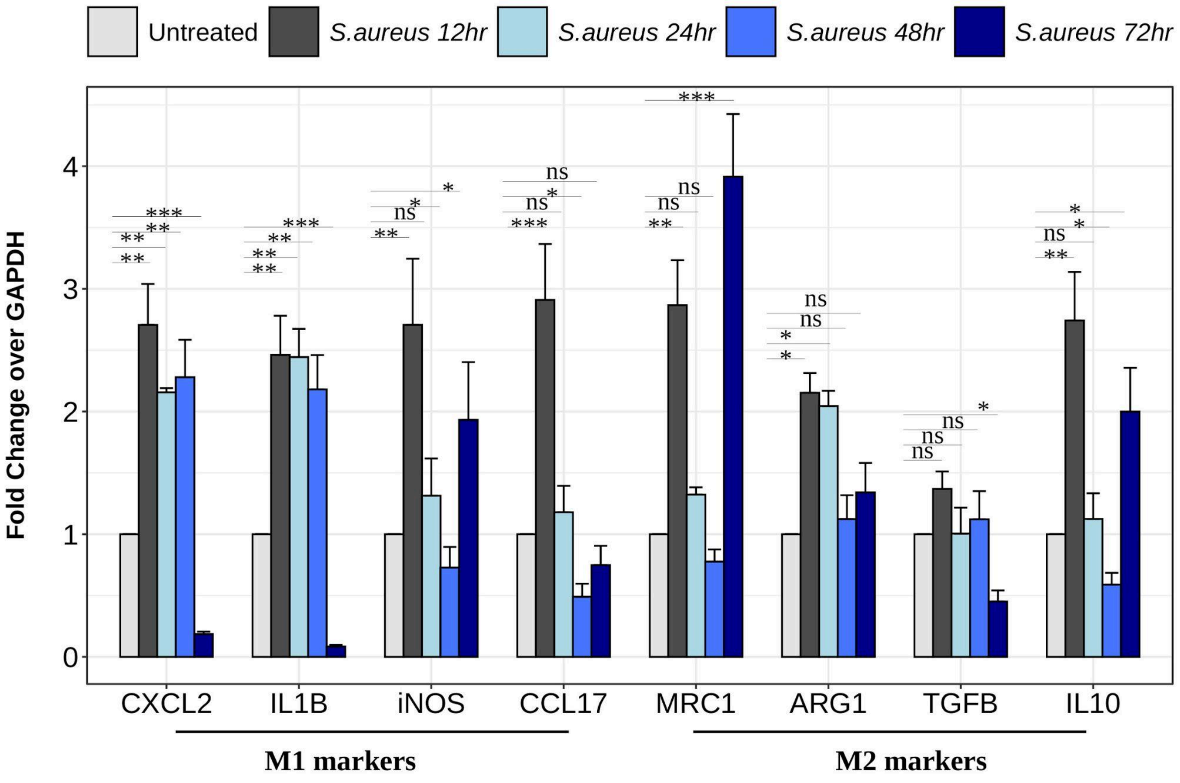


**Fig S10: Gene expression changes in M1 and M2 markers upon S.aureus infection at different time points.**

THP1 monocytes were treated with PMA (20ng/ml) for 16hr. Post 24hr, infection with S.aureus at 1:10MOI was given for mentioned time duration. Transcript levels of (A) M1 markers and (B) M2 markers were assessed. {Two-way ANOVA, $P > 0.05$ :ns, $P < 0.05$ :*, $P<0.01$ :**, $P<0.001$ :***, N=3 biological replicates}

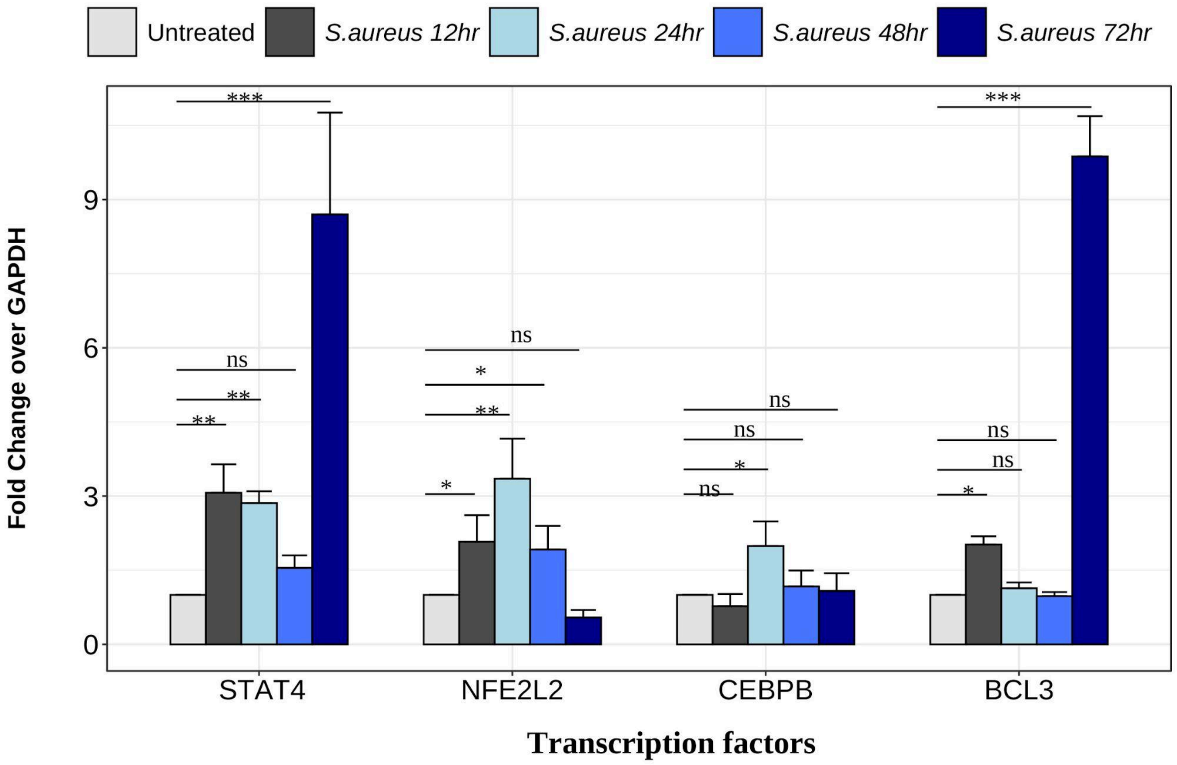


**Fig S11: Gene expression changes in STAT4 and NCB-set upon S.aureus infection at different time points.**

THP1 monocytes were treated with PMA (20ng/ml) for 16hr. Post 24hr, infection with S.aureus at 1:10MOI was given for mentioned time duration. Transcript levels of STAT4, NFE2L2, BCL3 adn CEBPB were assessed. {Two-way ANOVA, $P > 0.05$ :ns, $P < 0.05$ :*, $P<0.01$ :**, $P<0.001$ :***, N=3 biological replicates}.

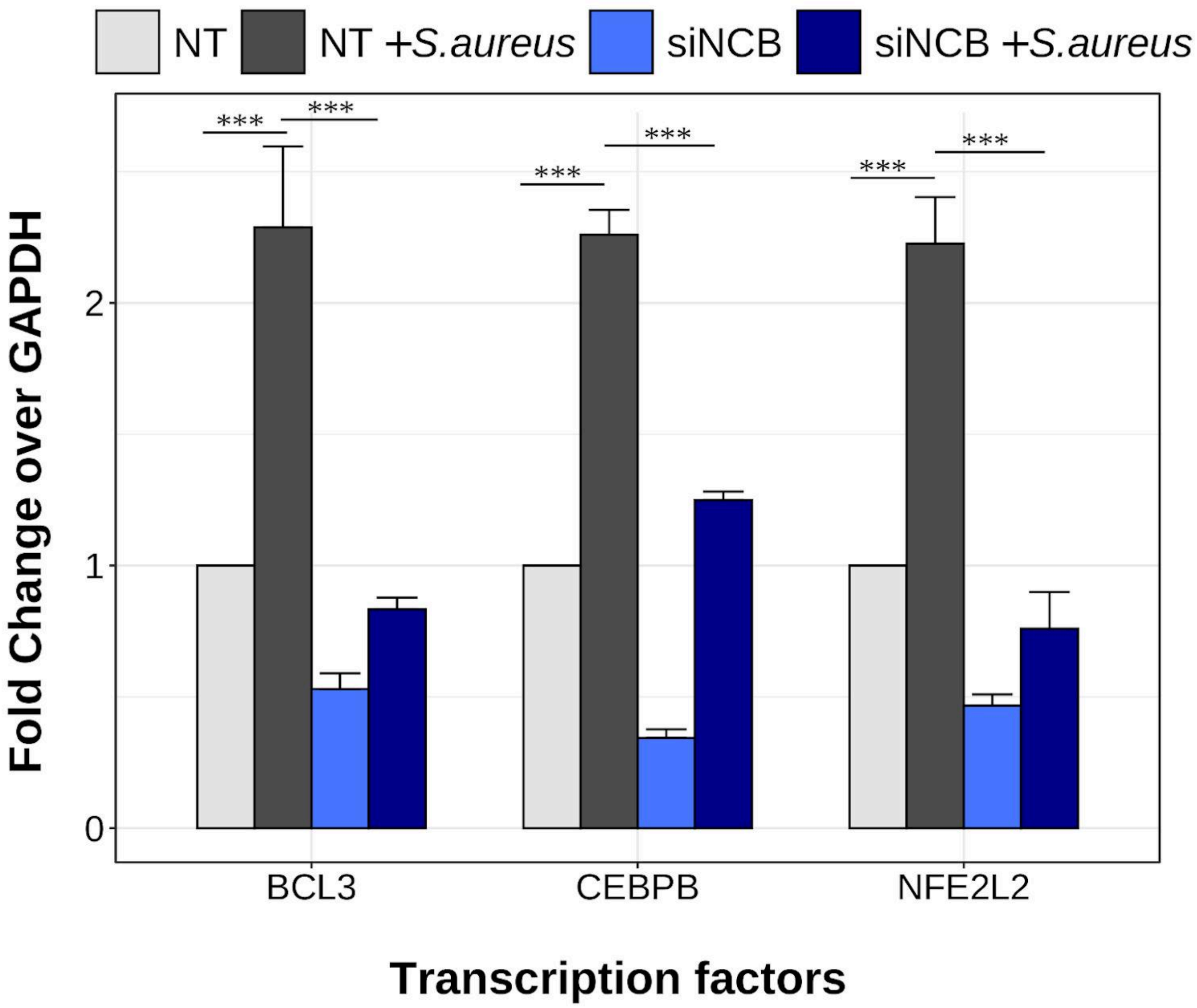


**Fig S12: NCB (NFE2L2, CEBPB, BCL3) siRNA validation in S.aureus data.**

THP1 monocytes were treated with PMA (20ng/ml) for 16hr. NFE2L2, CEBPB and BCL3 specific siRNAs were transfected. Post 24hr, S.aureus infection at 1:10 MOI was given for 24hr. Transcript levels of mentioned transcription factors were assessed. {Two-way ANOVA, **, P<0.001 :***, N=3, NT: non-targeting siRNA}

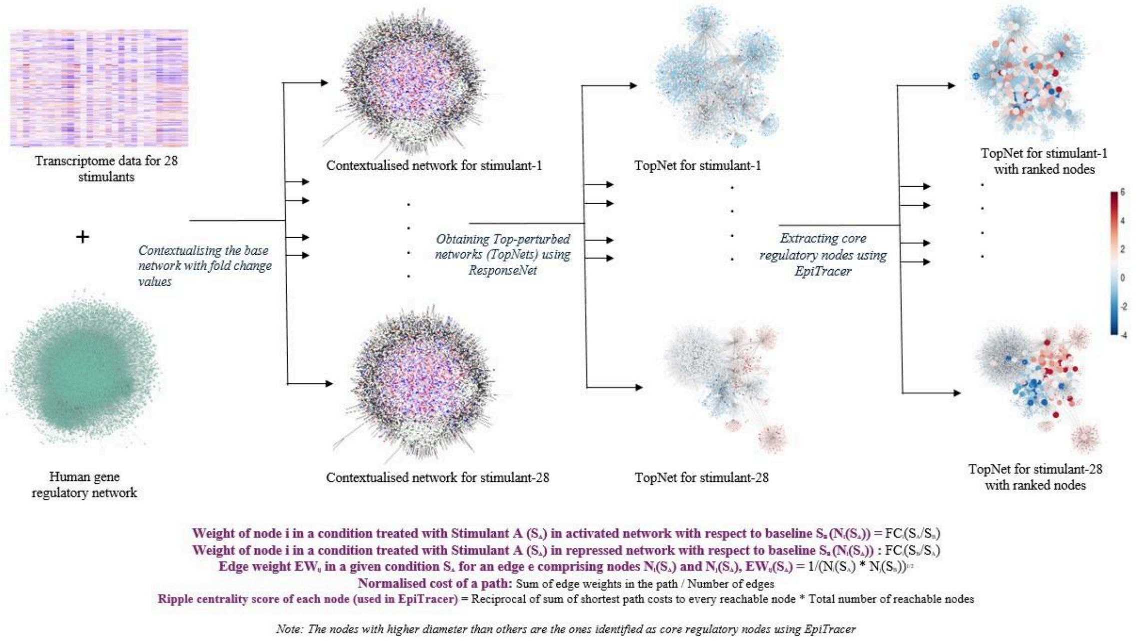


**Fig S13: Schematic diagram describing the network mining algorithm**

**Supplementary tables**

**Table S1.** ***In vitro stimulant data***. Details on the triggers and the end time point considered for the transcriptome analysis. (hrs – hours)

| **Stimulant(s)** | **Time Point** | **# Samples** |
|---|---|---|
| Baseline (Mϕ) () | 6hrs, 24hrs, 72 hrs | 3;3;7 |
| High Density Lipoprotein (HDL) | 6 hrs | 4 |
| Lauric Acid (LA) | 24 hrs | 6 |
| Linoleic Acid (LiA) | 24 hrs | 6 |
| Oleic Acid (OA) | 24 hrs | 9 |
| Palmitic Acid (PA) | 24 hrs | 9 |
| Stearic Acid (SA) | 24 hrs | 3 |
| Glucocorticoids (GC) | 72 hrs | 3 |
| Standard lipopolysaccharide (sLPS) | 72 hrs | 8 |
| Ultrapure Lipopolysaccharides (upLPS) | 72 hrs | 6 |
| Interferon beta (IFNβ) | 72 hrs | 3 |
| Interferon gamma (IFNγ) | 72 hrs | 7 |
| Interleukin-10 (IL10) | 72 hrs | 3 |
| Interleukin-13 (IL13) | 72 hrs | 3 |
| Interleukin-4 (IL4) | 72 hrs | 10 |
| Pam3CysSerLys4 (P3C) | 72 hrs | 3 |
| Prostaglandin $E_2$ (PGE2) | 72 hrs | 3 |
| P3C + PGE2 | 72 hrs | 3 |
| Tumour Necrosis Factor (TNF) | 72 hrs | 6 |
| sLPS+IC | 72 hrs | 3 |
| sLPS + IFNγ | 72 hrs | 3 |
| IFNγ +TNF | 72 hrs | 3 |
| TPP (TNF + PGE2 +P3C) | 72 hrs | 10 |
| TPP + IFNβ | 72 hrs | 3 |
| TPP + IFNβ + IFNγ | 72 hrs | 3 |
| TNF alpha + P3C | 72 hrs | 3 |

| | | |
|---|---|---|
| TNF alpha +PGE2 | 72 hrs | 3 |
| IL4 + upLPS | 72 hrs | 3 |
| upLPS +IC | 72 hrs | 3 |

**Table S2. The list of primers used for the qRT-PCT**

| Gene | Primer | Sequence |
|---|---|---|
| CEBPB | Forward | ATGTTCCTACGGGCTTGTTG |
| | Reverse | CCCAAAAGGCTTTGTAACCA |
| NFE2L2 | Forward | GCGACGGAAAGAGTATGAGC |
| | Reverse | GTTGGCAGATCCACTGGTTT |
| BCL3 | Forward | CCCTATACCCCATGATGTGC |
| | Reverse | GGTGTCTGCCGTAGGTTGTT |
| STAT4 | Forward | TACCTAATGCTTGGGCATCC |
| | Reverse | TCTGCCAGCATATGGAGTTG |
| IL1B | Forward | ATGATGGCTTATTACAGTGGCAA |
| | Reverse | GTCGGAGATTCGTAGCTGGA |
| IL10 | Forward | TGCCTTCAGCAGAGTGAAGA |
| | Reverse | TGGGTCTTGGTTCTCAGC |
| ARG1 | Forward | ACAGTTTGGCAATTGGAAGCA |
| | Reverse | CACCCAGATGACTCCAAGATCAG |
| MRC1 | Forward | GGGTTGCTATCACTCTCTATGC |
| | Reverse | TTTCTTGTCTGTTGCCGTAGTT |
| TGFB | Forward | CTAATGGTGGAAACCCACAACG |
| | Reverse | TATCGCCAGGAATTGTTGCTG |
| CXCL2 | Forward | CGCCCATGGTTAAGAAAATCA |
| | Reverse | CCTTCTGGTCAGTTGGATTTGC |
| iNOS | Forward | CAGCGGGATGACTTTCCAA |
| | Reverse | AGGCAAGATTTGGACCTGCA |
| SOCS3 | Forward | GTCACCCACAGCAAGTTT |
| | Reverse | CTGAGCGTGAAGAAGTGG |

**Table S3. The list of clusters, associated members, network details of regulatory cores and the number of TFs in the barcodes.** * corresponds to the clusters containing a single member. Nodes correspond to the number of genes and edges correspond to the number of interactions in the network.

| Cluster ID | Cluster members | Core Network statistics (Nodes ; | No. of Tfs in the Cluster centric transcription factor based barcode |
|---|---|---|---|

| | | Edges) | |
|---|---|---|---|
| Cluster 1 | P3C, P3C+PGE2, PGE2 | 2427; 2747 | 79 |
| Cluster 2* | IL4+upLPS | 4959; 6498 | 78 |
| Cluster 3 | IL13, IL4 | 1829; 1970 | 42 |
| Cluster 4 | GC, IL10 | 289; 302 | 5 |
| Cluster 5 | upLPS, upLPS+IC | 1576; 1698 | 43 |
| Cluster 6 | HDL, IFNβ | 901; 992 | 66 |
| Cluster 7 | TNFα, IFNγ+TNFα, sLPS+IFNγ | 364; 376 | 31 |
| Cluster 8 | TPP+IFNβ, TPP+IFNβ+IFNγ | 3103; 3432 | 113 |
| Cluster 9 | TNFα+P3C, TNFα+PGE2 | 3812; 4172 | 87 |
| Cluster 10 | OA, LiA, LA | 1922; 2059 | 30 |
| Cluster 11 | sLPS, sLPS+IC, TPP | 2028; 2145 | 52 |
| Cluster 12* | IFNγ | 5268; 6387 | 42 |
| Cluster 13 | PA, SA | 3482; 3945 | 68 |
